# The dual PPAR-α/δ agonist elafibranor attenuates TGF-β_1_-induced cardiac fibrosis through redox-metabolic and bioenergetic reprogramming in human cardiac models

**DOI:** 10.64898/2026.08.18.745425

**Authors:** Milena Paw, Lukas Minder, Andrea Laimbacher, Magdalena Czepiec, Sylwia Bobis-Wozowicz, Dawid Wnuk, Barbara Kutryb-Zając, Alicja Braczko, Michał Sarna, Patrycja Kaczara, Stefan Chłopicki, Zbigniew Madeja, Oliver Distler, Przemysław Błyszczuk, Jarosław Czyż, Gabriela Kania

## Abstract

**Background:** Cardiac fibrosis drives adverse myocardial remodelling through persistent fibroblast activation, ECM deposition, and impaired cardiac function. Current therapies offer limited protection against cardiac fibrosis progression. Elafibranor is a dual PPAR-α/δ agonist approved for the treatment of liver disease. However, its effects in human models of cardiac fibrosis remain insufficiently explored.

**Methods:** Elafibranor was evaluated in complementary human *in vitro* TGF-β_1_-induced cardiac fibrosis models: 2D primary fibroblasts, 3D fibroblast spheroids, spontaneously contracting 3D cardiac microtissues, and hiPSC-derived cardiomyocytes. Viability, apoptosis, fibroblast activation, ECM remodelling, mitochondrial respiration, nucleotide and NAD pools, calcium handling, contractility, and transcriptomic profiles were assessed.

**Results:** At non-cytotoxic concentrations, elafibranor attenuated TGF-β_1_-driven cardiac fibrosis responses. In 2D cardiac fibroblasts, it reduced myofibroblast differentiation, procollagen 1α1 secretion, and partially restored mitochondrial respiratory capacity. In 3D spheroids, it preserved viability, attenuated caspase-3/7 activation, and suppressed procollagen 1α1 release. In cardiac microtissues, elafibranor reduced ECM accumulation, shifted transcriptomic profiles toward redox-metabolic/cytoprotective pathways, altered adenine nucleotide and NAD pools, and partially recovered contraction parameters. In hiPSC-derived cardiomyocytes, elafibranor modulated calcium handling, contractility, and mitochondrial respiration.

**Conclusions:** Elafibranor mitigates TGF-β_1_-driven cardiac fibrosis by suppressing fibroblast activation and ECM remodelling while promoting adaptive metabolic, redox, and bioenergetic responses, supporting balanced PPAR-α/δ activation as a potential therapeutic strategy for cardiac fibrosis.

**Graphical abstract:** 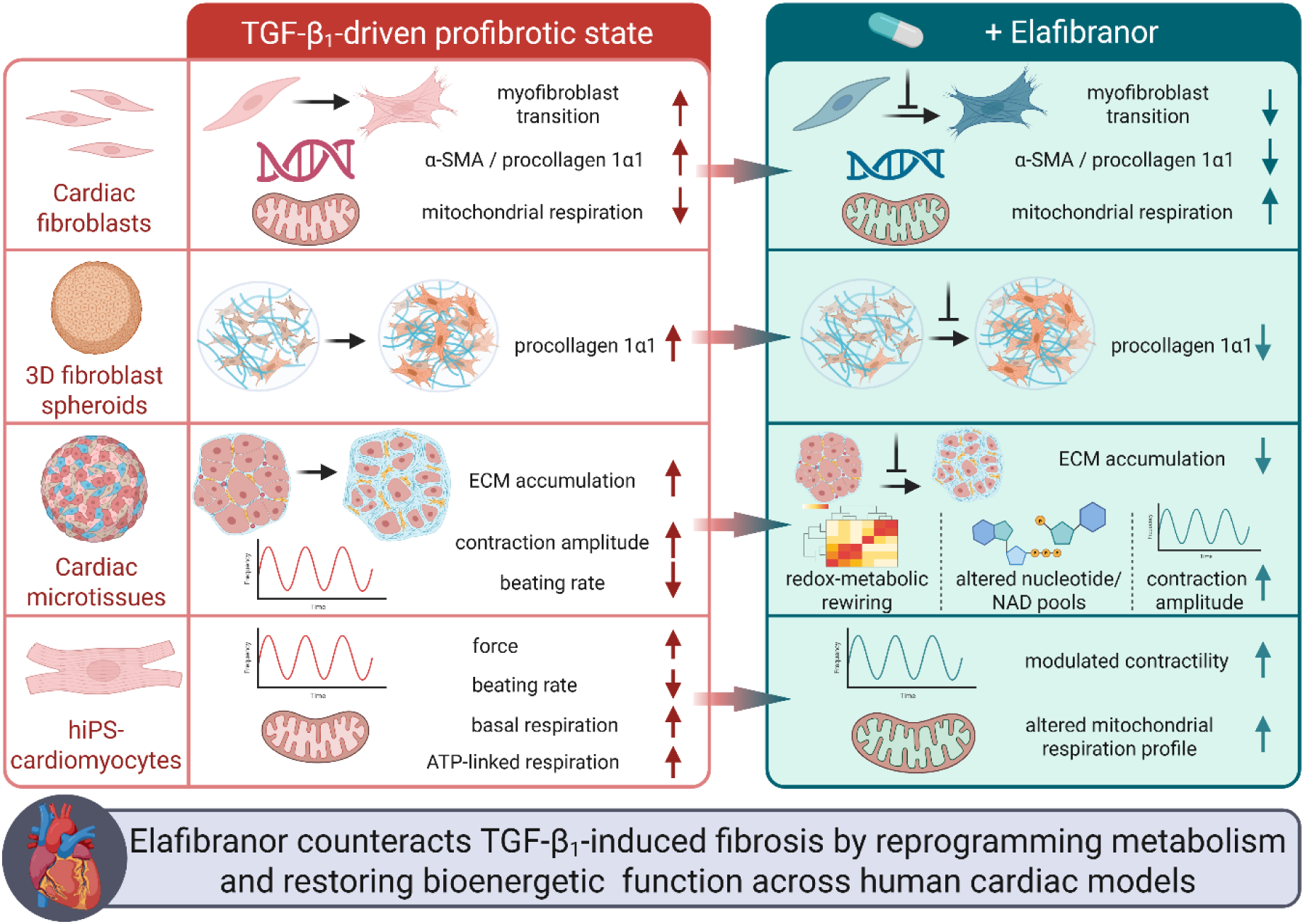

## 1. Background

Cardiovascular diseases (CVDs) remain the leading cause of mortality worldwide. Recent Global Burden of Disease estimates indicate approximately 20.5 million CVD deaths in 2021 and more than 600 million affected individuals [1–3]. Despite advances in prevention and acute care, chronic cardiac conditions continue to rise with population aging and persistent cardiometabolic risk factors [1]. Across heart diseases, cardiac fibrosis is central to pathological myocardial remodeling and progression toward heart failure [4,5]. It involves excessive extracellular matrix (ECM) deposition and reorganization, causing myocardial stiffening, disrupted architecture, impaired mechanical/electrical coupling, functional decline, and increased arrhythmogenic susceptibility [4,6]. Rather than a passive consequence of injury, cardiac fibrosis is actively regulated by cellular, mechanical, and metabolic signals [7,8].

Cardiac fibroblasts are the principal matrix-producing cells and the dominant source of myofibroblasts during remodeling [9]. Single-cell transcriptomic and spatial genomic studies revealed substantial fibroblast heterogeneity, with distinct activation states in healthy and diseased myocardium [10,11]. Fibroblast activation is driven by transforming growth factor-β, angiotensin II, endothelin-1, inflammatory cytokines, and matrix stiffness, which reinforce cytoskeletal tension and mechanotransduction, stabilize the myofibroblastic phenotype, and sustain ECM overproduction [4,12–16]. If unresolved, persistent myofibroblast activity promotes ECM accumulation and myocardial stiffening, features closely linked to impaired diastolic function [17]. Although renin-angiotensin-aldosterone system (RAAS) inhibition and mineralocorticoid receptor antagonist’s slow fibrotic progression and reduce profibrotic signaling, they do not reverse established fibrotic tissue in the human heart, highlighting an unmet therapeutic need [4].

This supports targeting molecular programs regulating fibroblast activation and persistence, including peroxisome proliferator-activated receptors (PPARs). PPARs are ligand-activated nuclear receptors comprising PPAR-α, PPAR-β/δ, and PPAR-γ, which regulate lipid metabolism, inflammation, and cellular differentiation in a context-dependent manner [18,19]. Clinically, PPAR-α agonists treat dyslipidemia and PPAR-γ agonists treat type 2 diabetes, whereas no PPAR-δ-selective agent is established for routine clinical use [20]. In cardiac models, PPAR signaling modulates remodeling in an isoform- and context-dependent manner: PPAR-δ activation promoted early infarct healing by enhancing angiogenesis, recruitment of bone marrow-derived mesenchymal cells, TGF-β_2_ expression, fibroblast-to-myofibroblast differentiation, collagen deposition, and matrix remodeling after myocardial infarction, whereas PPAR-α activation preserved myocardial energetics by maintaining fatty acid oxidation and high-energy phosphate levels while attenuating fibrosis-related gene induction in pressure-overload heart failure [21,22].

Elafibranor (GFT505) is a dual PPAR-α/δ agonist originally developed for metabolic liver disease. In preclinical nonalcoholic steatohepatitis (NASH) models, it reduced inflammatory and profibrotic gene expression and attenuated fibrosis progression [23]. In the phase IIb GOLDEN-505 trial, elafibranor showed biological activity in NASH, including favorable metabolic and inflammatory effects; although the primary endpoint was not met, post-hoc analyses using modified criteria indicated resolution of steatohepatitis without worsening of fibrosis in selected subgroups, particularly patients with more advanced disease [24]. More recently, elafibranor improved diastolic dysfunction in a preclinical model of NASH-associated heart failure with preserved ejection fraction (HFpEF), supporting its relevance to cardiometabolic cardiac remodeling [25]. These findings support investigation of balanced PPAR-α/δ activation in cardiac fibroblast-dependent remodeling.

However, the impact of elafibranor on cardiac fibrosis remains insufficiently explored. Thus, we investigated its anti-fibrotic potential in human-relevant 2D cultures and 3D cardiac microtissues (MTs) [26,27], assessing fibroblast activation, ECM remodeling, and tissue-level functional properties.

## 2. Materials and methods

### 2.1. Cell cultures and MTs formation

Fetal and adult human cardiac fibroblasts (hCFs; Cell Applications/Merck, cat. no. 306-05F and 306-05A, respectively) were cultured under standard conditions (37 °C, 5% CO₂) in Dulbecco’s Modified Eagle Medium with high glucose (DMEM-HG; Sigma-Aldrich/Merck), supplemented with 10% fetal bovine serum (FBS; Gibco) and a penicillin/streptomycin cocktail (Gibco). Experiments based on the 2D cultures of hCFs were performed using cells between 6th and 15^th^ passages. Cells were seeded at a density of 5 000 cells/cm². After 24 hours, the culture medium was replaced with Maintenance Medium (MM; DMEM-HG supplemented with 2% FBS, 50 μM phenylephrine hydrochloride (Sigma-Aldrich/Merck), 0.3 μM L-ascorbic acid (Sigma-Aldrich/Merck), and 50 U/mL penicillin/streptomycin (Gibco) without or with elafibranor (Cayman Chemical; 0 - 60 μM). Recombinant human TGF-β_1_ (PeproTech; 10 ng/ml) was used to the induction of fibrotic response. Cells were cultured under these conditions for 4 days. For spheroid formation, hCFs were suspended in MM and seeded into 96-well GravityTRAP plates (InSphero, Schlieren, Switzerland). 3D hCFs cultures were incubated under standard conditions (37 °C, 5% CO₂) in a tilted position for 2 days to facilitate self-assembly and then the medium was replaced by fresh MM without or with tested compounds. Spheroids were maintained for 10 days, with medium changes performed every 2 days.

Human induced pluripotent stem cells-derived cardiomyocytes (hiPSC-CMs) were obtained either from FujiFilm Cellular Dynamics (iCell® Cardiomyocytes2, cat. #01434) or generated from the episomal hiPSC line (Gibco, A18945) as previously described [28]. For 2D experiments, hiPSC-CMs were seeded on Geltrex-coated plates (LDEV-free, reduced growth factor basement membrane matrix; Gibco) in RPMI-1640 medium supplemented with 2% B27 with insulin (Gibco). Metabolic selection using 4 mM sodium lactate (Sigma-Aldrich/Merck) in glucose-free DMEM (Gibco) was performed to obtain purified CM cultures. Cells were stimulated with elafibranor (15 μM), without or with TGF-β_1_ (10 ng/ml), and cultured under standard conditions (37 °C, 5% CO₂) for 24 hours.

To assembly MTs, co-cultures of hiPSC-CMs and hCFs were used. Cells were mixed at a 4:1 ratio (hiPSC-CMs:CFs, 5000 cells/spheroid), suspended in 80 μl of MM and seeded into each well of 96-well GravityTRAP plates (InSphero, Schlieren, Switzerland). Cultures were incubated under standard conditions (37 °C, 5% CO₂) in a tilted position for 2 days to facilitate self-assembly. Then, the medium was replaced with fresh MM containing elafibranor (0 - 60 μM) in the absence or presence of TGF-β_1_ (20 ng/ml). MTs were cultured for 10 days, with medium changes performed every 2 days.

### 2.2. Evaluation of contractile function of MTs and hiPSC-CMs

Contractile activity of hiPSC-CMs was measured using an integrated AFM-fluorescence microscopy setup consisting of a BioScope Catalyst atomic force microscope coupled to a Zeiss inverted microscope and an ORCA-Flash4.0 LT3 digital CMOS camera. Prior to imaging, cells were loaded with Fluo-4 (Invitrogen) for 30 min to enable simultaneous monitoring of calcium transients. Measurements were carried out at 37 °C after positioning selected cells within the optical field of the AFM system. For each experimental group, 10 individual cells were analyzed. AFM recordings were obtained using an MLCT-Bio probe with a nominal spring constant of 0.01 N/m, which was brought into contact with each cell before data acquisition. Force and fluorescence signals were recorded simultaneously using a custom acquisition script, with identical camera and imaging settings maintained for all fluorescence measurements. Raw AFM traces were converted into force values using calibration files and in-house software. Probe calibration was performed both before and after each measurement session, and the averaged calibration values were used for subsequent calculation of beating rate and contraction force. Calcium flux was analyzed from fluorescence recordings using a dedicated ImageJ plugin.

MT contractility was assessed using the Axio Observer Z1 microscope (Zeiss, Hombrechtikon, Switzerland) and ZEN software. Captured images were converted into videos using Fiji (ImageJ) software and a custom-made macro. Contractile properties of the MTs were then analyzed using Fiji and the MUSCLEMOTION macro [29].

### 2.3. Cell viability and metabolic activity assays

Cell metabolic activity and viability were evaluated after treatment with increasing concentrations of elafibranor (0-60 μM), applied either alone or together with TGF-β₁. TGF-β₁ was used at 10 ng/ml in 2D cultures and at 20 ng/ml in 3D spheroid cultures. Metabolic activity was assessed using two independent colorimetric/fluorometric assays: PrestoBlue™ HS Cell Viability Reagent (Invitrogen) and MTT. Both assays were performed according to the manufacturers’ protocols. For the PrestoBlue assay, the reagent was diluted 1:10 in fresh maintenance medium, added directly to wells containing cells or spheroids, and incubated for 4 h at 37 °C in 5% CO₂. For the MTT assay, MTT solution (5 mg/ml; Sigma-Aldrich/Merck) was added to conditioned medium to obtain a final concentration of 0.5 mg/ml and incubated with the cultures for 4 h under standard culture conditions. After incubation, the medium was removed and formazan crystals were dissolved in 100 μl of isopropanol per well. Resorufin fluorescence was read at 590 nm, while solubilized formazan absorbance was measured at 570 nm using either a Synergy microplate reader with Gen5 software (BioTek) or a Multiskan FC reader (Thermo Fisher Scientific).

Cell membrane integrity and viability were further analyzed using fluorescein diacetate/ethidium bromide (FDA/EtBr; Sigma-Aldrich/Merck) staining. After treatment, cells were detached with trypsin, resuspended in phosphate-buffered saline (PBS) containing FDA/EtBr staining solution, and immediately examined by fluorescence microscopy. Viable cells were identified as FDA⁺/EtBr⁻, whereas EtBr⁺ cells were classified as non-viable. Cell counting was performed using a Leica DMI6000B fluorescence microscope equipped with LAS X software (Leica Microsystems GmbH, Wetzlar, Germany). Viability was calculated as the percentage of viable cells in each treated condition (7.5-60 μM elafibranor) relative to untreated controls (0 μM).

### 2.4. Caspase 3/7 activity assay

MTs were transferred into fresh medium in white-walled 96-well plates, and an equal volume of Caspase-Glo® 3/7 reagent (Promega, Dübendorf, Switzerland) was added to each well. After gentle mixing, the plates were incubated at room temperature for 3 hours, protected from light. Luminescence was then measured using a Synergy microplate reader (BioTek, Winooski, Vermont, USA) and analyzed with Gen5 software.

### 2.5. á-SMA and procollagen 1α1 quantification by ELISA

After 4 days of culture under the 2D conditions described above, hCFs were processed for α-SMA detection. Cells were first fixed and permeabilized with pre-cooled methanol, followed by blocking for 1 h in PBS containing 1% BSA and 0.1% Tween-20. The cultures were then incubated overnight at 4 °C with a mouse monoclonal antibody directed against α-SMA (Sigma-Aldrich/Merck), prepared in 1% BSA/PBS. After washing the cells three times, HRP-labeled goat anti-mouse secondary antibodies were added for 1 h at room temperature using the same antibody diluent. Signal development was performed with tetramethylbenzidine (TMB; Sigma-Aldrich/Merck), and the reaction was stopped with 1N HCl. The resulting absorbance was measured at 450 nm using a Synergy plate reader (BioTek, Winooski, VT, USA) or a Multiskan FC reader (Thermo Fisher Scientific).

For procollagen 1α1 quantification, conditioned media were harvested from 2D hCF cultures after 4 days and from individual 3D MTs after 10 days of culture. Samples were kept at -80 °C until measurement. Procollagen 1α1 concentration was determined with the Human Procollagen 1α1 DuoSet ELISA kit (RCD Systems, Abingdon, UK), following the manufacturer’s instructions. Absorbance was read at 450 nm using a Synergy microplate reader (BioTek, Winooski, VT, USA), and results were analyzed with Gen5 software.

### 2.6. Immunofluorescence and histochemical staining and image analysis

For immunofluorescence staining, cells were plated on glass coverslips placed in 12-well plates and maintained for 4 days under the culture conditions described above. After treatment, cells were fixed and stained following the previously described protocol [30]. A mouse monoclonal IgG antibody against α-SMA (Sigma-Aldrich/Merck) was used as the primary antibody, followed by the appropriate Alexa Fluor 488-conjugated secondary antibodies (Life Technologies, Thermo Fisher Scientific, Waltham, MA, USA). Cell nuclei were counterstained with Hoechst 33258 (1 µg/ml; Sigma-Aldrich/Merck). Fluorescence images were captured using a Leica DMI6000B microscope operated with LAS X software version 3.7.4 (Leica Microsystems GmbH, Wetzlar, Germany).

For immunohistochemical analysis of 3D cultures, 25-30 cardiac MTs were pooled per sample, fixed overnight in 4% paraformaldehyde prepared in PBS, and embedded in 1% agarose before routine paraffin processing. Samples were dehydrated through graded ethanol solutions, cleared in xylene, and embedded in paraffin at 56 °C. Sections of 2 µm thickness were prepared on Superfrost Plus slides and dried overnight at 58 °C. Before staining, sections were deparaffinized in xylene, rehydrated through decreasing ethanol concentrations, rinsed in distilled water, and subjected to heat-induced antigen retrieval using either Tris-EDTA buffer at pH 9 or citrate buffer at pH 6. Non-specific binding was blocked with 10% goat serum (Vector Laboratories). Sections were then incubated with primary antibodies against α-SMA (Abcam ab132575, clone E184; 1:2000), periostin (Abcam ab14041; 1:2000), fibronectin (Abcam ab2413; 1:2000), and troponin T (Invitrogen MA4-12960; 1:20000). Immunoreactivity was visualized with the Bond Polymer HRP Refine Detection Kit (Leica), and nuclei were counterstained with haematoxylin. Collagen deposition was evaluated using Masson’s Trichrome staining. Stained sections were imaged on a Leica DMi8 microscope equipped with CoolLED pE-4000 illumination and AFC autofocus stabilization. Quantification of immunoreactive signal and nuclear counterstaining was performed in Fiji. Signal intensity was normalized to the number of counterstained nuclei and expressed as arbitrary units (AU).

### 2.7. Intracellular nucleotide quantification

For intracellular nucleotide and NAD analysis, MTs were collected in 100 μl of 80% methanol and kept at -80 °C until processing. Samples were prepared and analyzed by UHPLC-DAD as described previously [31,32]. Measurements were performed using a Nexera LC-40 UHPLC system equipped with an SPD-M30A diode array detector and a high-sensitivity 85-mm optical path flow cell (Shimadzu, Japan). Based on the quantified nucleotide levels, ATP/AMP and ATP/ADP ratios, the total adenine nucleotide pool (TAN; ATP + ADP + AMP), and NAD content were determined. Adenylate energy charge (AEC) was calculated as (ATP + 0.5 ADP)/(ATP + ADP + AMP), according to Atkinson’s definition of the adenylate pool energy charge [33].

### 2.8. Mitochondrial respiration analysis

For extracellular flux analysis, hCFs and hiPSC-CMs were plated in Seahorse XFe96 cell culture microplates at densities of 5,000 and 15,000 cells per well, respectively, using the appropriate complete culture medium for each cell type. After 24 h, the medium was exchanged for fresh medium containing elafibranor, either alone or in combination with TGF-β₁, and cells were maintained for an additional 4 days. Mitochondrial respiration was assessed using a Seahorse analyzer (Seahorse Bioscience, North Billerica, MA, USA) and the Agilent Seahorse XF Cell Mito Stress Test Kit. One day before the assay, the Seahorse XF sensor cartridge was hydrated overnight at 37°C in a non-CO₂ incubator according to the manufacturer’s instructions. On the day of the assay, culture medium was replaced with bicarbonate-free Agilent Seahorse XF Base Medium Minimal DMEM or DMEM (D5030, Sigma-Aldrich), supplemented with 10 mM glucose, 2 mM L-glutamine, and 1 mM sodium pyruvate, adjusted to pH 7.4. Cells were washed twice with the appropriate assay medium and equilibrated for 45 min at 37 °C in a CO₂-free incubator. Concentrations of mitochondrial modulators were optimized separately for hCFs and hiPSC-CMs in preliminary experiments. During the mitochondrial stress test, compounds were injected sequentially as follows: oligomycin, an ATP synthase inhibitor, at 1 μg/ml; FCCP, a mitochondrial uncoupler, at 9 μM for hCFs or 2 μM for hiPSC-CMs; and rotenone plus antimycin A, inhibitors of complexes I and III, at 0.5 μM each. Basal respiration, ATP-linked respiration, maximal respiration, and spare respiratory capacity were derived from OCR traces using Seahorse Wave software. OCR data were normalized either to cell number or to total protein content, as specified. Each condition was analyzed using multiple technical replicates across three independent biological experiments.

### 2.9. RNA sequencing and data analysis

RNA isolation, library preparation, sequencing, and primary data processing were performed by Lexogen. The experiment included 12 samples representing four experimental conditions, with three biological replicates per condition. Libraries were prepared using the Lexogen LUTHOR workflow for low-input 3′ mRNA-seq and included unique molecular identifiers (UMIs) for gene-level deduplication. Sequencing was performed in paired-end mode, and data were processed with Lexogen DAP v1.7.7. Across samples, the number of input reads ranged from 3.34 to 7.66 million, with a median of 4.92 million reads per sample; the corresponding number of uniquely mapped reads ranged from 3.08 to 7.08 million, with a median of 4.55 million. Reads were aligned with STAR to the human GRCh38.94 reference supplemented with ERCC and SIRV spike-in sequences. Alignment quality was high across all samples, with uniquely mapped reads ranging from 92.38% to 93.09%, and the average post-trimming read length was 93-95 bp. UMI-based deduplication was performed at the gene level before downstream gene-level quantification. Differential expression analysis was performed with DESeq2, and genes were considered differentially expressed using a conservative threshold of |log₂ fold change| ≥ 1 and Benjamini-Hochberg-adjusted p-value/FDR < 0.05. For pathway-level analysis, preranked GSEA was performed with fgsea using gene lists ranked by the DESeq2 Wald statistic. Hallmark, Reactome, and Gene Ontology Biological Process gene sets were obtained from msigdbr, with gene sets containing 15-500 genes tested and FDR < 0.05 considered significant. Reactome GSEA was used for the main mechanistic interpretation, while Hallmark and GO Biological Process outputs were used as complementary pathway-level analyses. As a supplementary DEG-based approach, Reactome over-representation analysis was performed separately for upregulated and downregulated DEGs using clusterProfiler::enricher, with all genes detected in the corresponding DESeq2 result table used as the background universe. Reactome gene sets containing 10-500 genes were tested, and FDR < 0.05 was used as the significance threshold. Downstream analyses and visualization were performed in R using custom scripts. RNA-seq data generated in this study will be deposited in the Gene Expression Omnibus, and accession numbers will be provided prior to publication.

### 2.10. Statistical analysis

Each experiment were performed in at least three replicates. Samples or data points were excluded only when predefined technical quality criteria were not met, including failed acquisition, poor image quality or damaged spheroids. All quantitative data are presented as mean ± standard deviation (SD). The normality of data distribution was assessed using the Shapiro-Wilk test. Depending on the distribution, statistical significance was determined using either the nonparametric Kruskal-Wallis test followed by Dunn’s multiple comparisons post hoc test, or one-way analysis of variance (ANOVA) with Tukey’s multiple comparisons post hoc test. Individual p-values are indicated in each graph. All statistical analyses were performed using GraphPad Prism version 10.4.2.

## 3. Results

### 3.1. Elafibranor attenuates the TGF-β₁-induced myofibroblastic transition of human cardiac fibroblasts in 2D cultures

To assess the cytotoxic and pro-apoptotic effects of elafibranor, human cardiac fibroblasts (hCFs) were cultured in 2D and treated with increasing concentrations of elafibranor (7.5-60 μM), either in the absence or presence of TGF-β_1_. Phase-contrast microscopy showed a typical spindle-like fibroblast morphology in control cells, whereas TGF-β_1_ induced cell flattening and an increased cell surface area. Co-treatment with elafibranor at concentrations ≥15 μM partially reversed these morphological changes, restoring a more spindle-like fibroblast phenotype (Figure 1A). Low concentrations of elafibranor (7.5-30 μM) modestly increased the MTT signal, whereas higher concentrations (≥45 μM) significantly reduced metabolic activity in both untreated and TGF-β_1_-stimulated hCFs (Figure 1B-C). These findings were supported by FDA/EtBr staining, which confirmed increased cell death at concentrations ≥45 μM irrespective of TGF-β_1_ treatment (Figure 1D). Consistently, bright-field images indicated reduced cell density at the highest concentration tested, particularly at 60 μM. Caspase-3/7 activity was significantly increased at elafibranor concentrations ≥45 μM in both control and TGF-β_1_-treated hCFs, indicating apoptosis induction (Figure 1E). Collectively, these data show that elafibranor is non-cytotoxic at low concentrations (7.5-30 μM), whereas higher concentrations (≥45 μM) induce cytotoxic and pro-apoptotic effects in hCFs independently of TGF-β_1_ stimulation.

**Figure 1.**
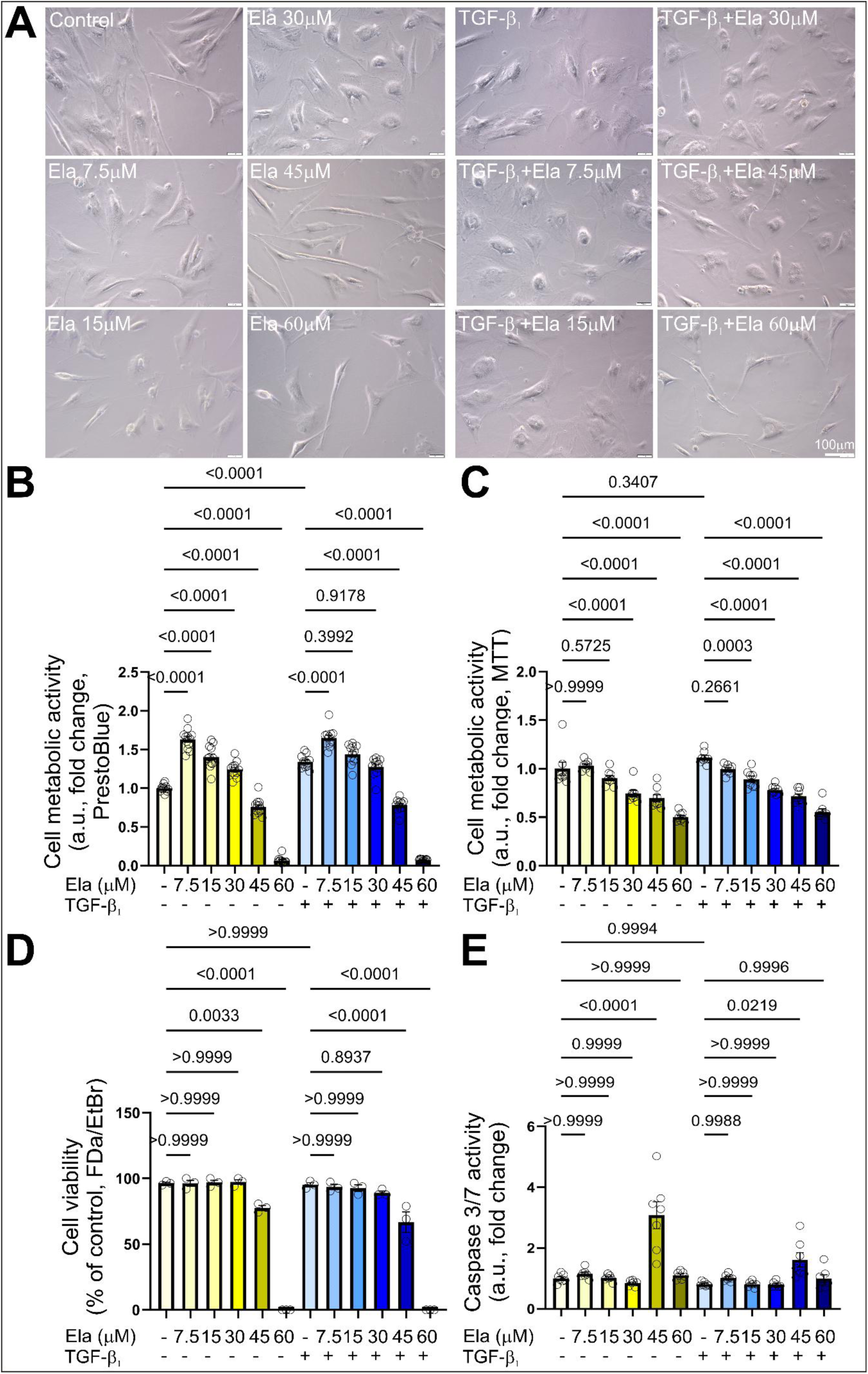
Low-dose elafibranor preserves hCF viability, whereas higher concentrations induce cytotoxic and pro-apoptotic effects. Cells were cultured n 2D standard conditions in the increasing concentrations of elafibranor (7.5, 15, 30, 45 and 60 μM) without or with TGF-β_1_ (10 ng/ml) for 4 days. **(A)** Representative phase-contrast images of cell morphology. Scale bar: 100 μm. Cell metabolic activity was determined using **(B)** PrestoBlue and **(C)** MTT assays. Cell viability was determined using **(D)** FDA/EtBr assay. **(E)** Quantification of caspase-3/7 activity in cells treated with elafibranor without or with TGF-β_1_. Data are presented as mean ± SD. Statistical significance was determined by one-way ANOVA followed by Tukey’s multiple comparisons test. p-values are indicated in each graph.

To investigate the antifibrotic potential of elafibranor, we assessed fibroblast-to-myofibroblast differentiation in hCFs by measuring α-smooth muscle actin (α-SMA) expression and secreted procollagen 1α1 levels following treatment with increasing concentrations of elafibranor (7.5-60 μM), in the absence or presence of TGF-β_1_. TGF-β_1_ significantly increased α-SMA expression and procollagen 1α1 secretion compared with untreated controls, and these effects were attenuated by elafibranor in a dose-dependent manner, with pronounced inhibition at concentrations ≥30 μM (Figure 2A-B). Since 15 μM elafibranor was selected as the non-cytotoxic working concentration for subsequent experiments, this concentration was used for immunofluorescence validation of α-SMA-positive stress fiber formation. Consistently, TGF-β_1_-induced formation of α-SMA-positive stress fibers was reduced upon co-treatment with 15 μM elafibranor (Figure 2C), resulting in a significant decrease in the proportion of α-SMA-positive myofibroblasts under standard 2D culture conditions (Figure 2D). Given that TGF-β_1_-driven fibroblast differentiation is associated with metabolic reprogramming, we next examined the effects of elafibranor at the same non-cytotoxic working concentration of 15 μM on cellular metabolism using Seahorse XF Mito Stress Test analysis. While TGF-β₁ did not affect basal respiration, it significantly reduced maximal respiration and spare respiratory capacity, accompanied by increased ATP-linked respiration (Figure 2E-I). Co-treatment with elafibranor at 15 μM partially improved basal respiration and fully reversed the TGF-β₁-induced impairments in maximal respiration and spare respiratory capacity to control levels (Figure 2F-H), without significantly altering ATP-linked respiration (Figure 2I). Together, these data demonstrate that elafibranor at 15 μM modulates cellular energy metabolism and attenuates TGF-β₁-induced myofibroblast differentiation in hCFs without inducing cytotoxic or pro-apoptotic effects.

**Figure 2.**
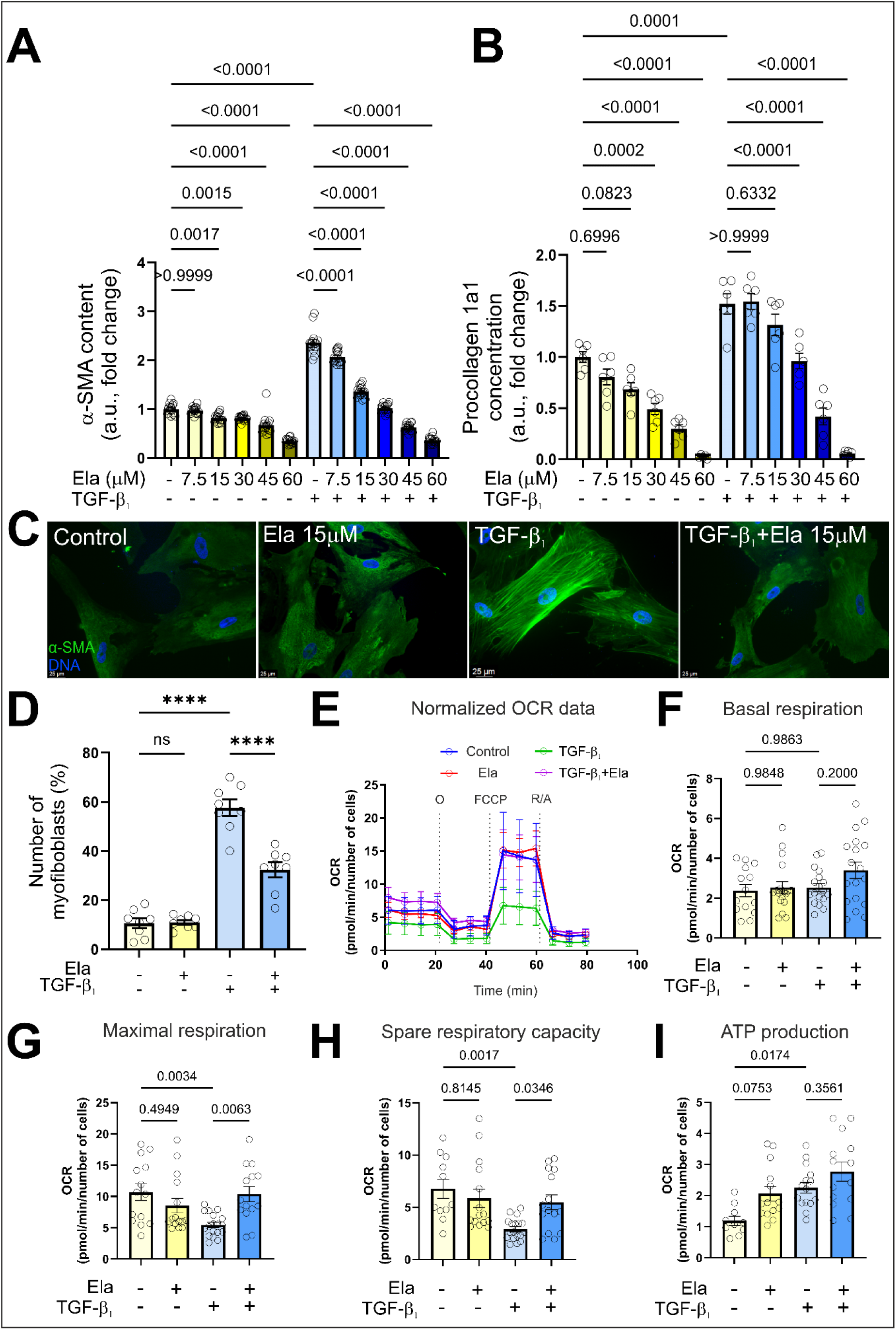
Elafibranor suppresses TGF-β_1_-induced myofibroblast transition and modulates mitochondrial respiration in hCFs. Cells were cultured in 2D standard conditions in the increasing concentrations of elafibranor (7.5, 15, 30, 45 and 60 μM) without or with TGF-β_1_ (10 ng/ml) for 4 days. **(A)** α-SMA protein levels in cultured hCFs was determined using in-cell ELISA assay. **(B)** Procollagen 1α1 concentration was determined in the conditioned medium from same hCF cultures using ELISA assay. **(C)** Representative images of cells immunostained for α-SMA (green) and DNA (blue). Scale bar: 25 μm. **(D)** Efficiency of fibroblast-myofibroblast differentiation was determined and presented as a percentage of α-SMA-positive myofibroblasts. **(E)** Trajectories and oxygen consumption rate (OCR) values were normalized to cell number for all tested conditions. Cellular bioenergetic changes was presented as **(F)** basal respiration, **(G)** maximal respiration, **(H)** spare respiratory capacity and **(I)** ATP-linked respiration. Data are presented as mean ± SD (n = biological replicates). Statistical significance was determined by one-way ANOVA followed by Tukey’s multiple comparisons test. p-values are indicated in each graph.

### 3.2. Elafibranor preserves viability and attenuates TGF-β_1_-induced procollagen 1α1 secretion in 3D hCF spheroids

To investigate the effects of elafibranor on TGF-β_1_-induced procollagen 1α1 secretion in a three-dimensional fibroblast culture system, we used hCF spheroids as a model of fibroblast-associated profibrotic activation. Spheroids were treated with increasing concentrations of elafibranor (7.5-60 μM), without or with TGF-β₁ stimulation for 10 days. Representative phase-contrast images revealed that fibroblast spheroids preserved their compact and spherical morphology in all elafibranor concentrations, irrespective of TGF-β₁ stimulation (Figure 3A). Cell viability was not significantly affected by any of the tested elafibranor concentrations, regardless of TGF-β₁ co-treatment, suggesting a lack of cytotoxic effects of the compound on 3D hCF cultures (Figure 3B). Furthermore, elafibranor administered alone did not affect caspase-3/7 activation. Although TGF-β₁ stimulation significantly increased caspase-3/7 activity, elafibranor at all tested concentrations restored it to baseline (control) levels, suggesting a robust anti-apoptotic effect (Figure 3C). Similarly, TGF-β₁-induced upregulation of procollagen 1α1 was significantly attenuated by elafibranor in a concentration-dependent manner (Figure 3D). Together, these findings indicate that elafibranor demonstrates a lack of cytotoxic and pro-apoptotic effects on hCF 3D spheroids and effectively suppresses profibrotic marker secretion in a concentration-dependent manner, supporting its therapeutic potential in cardiac fibrosis.

**Figure 3.**
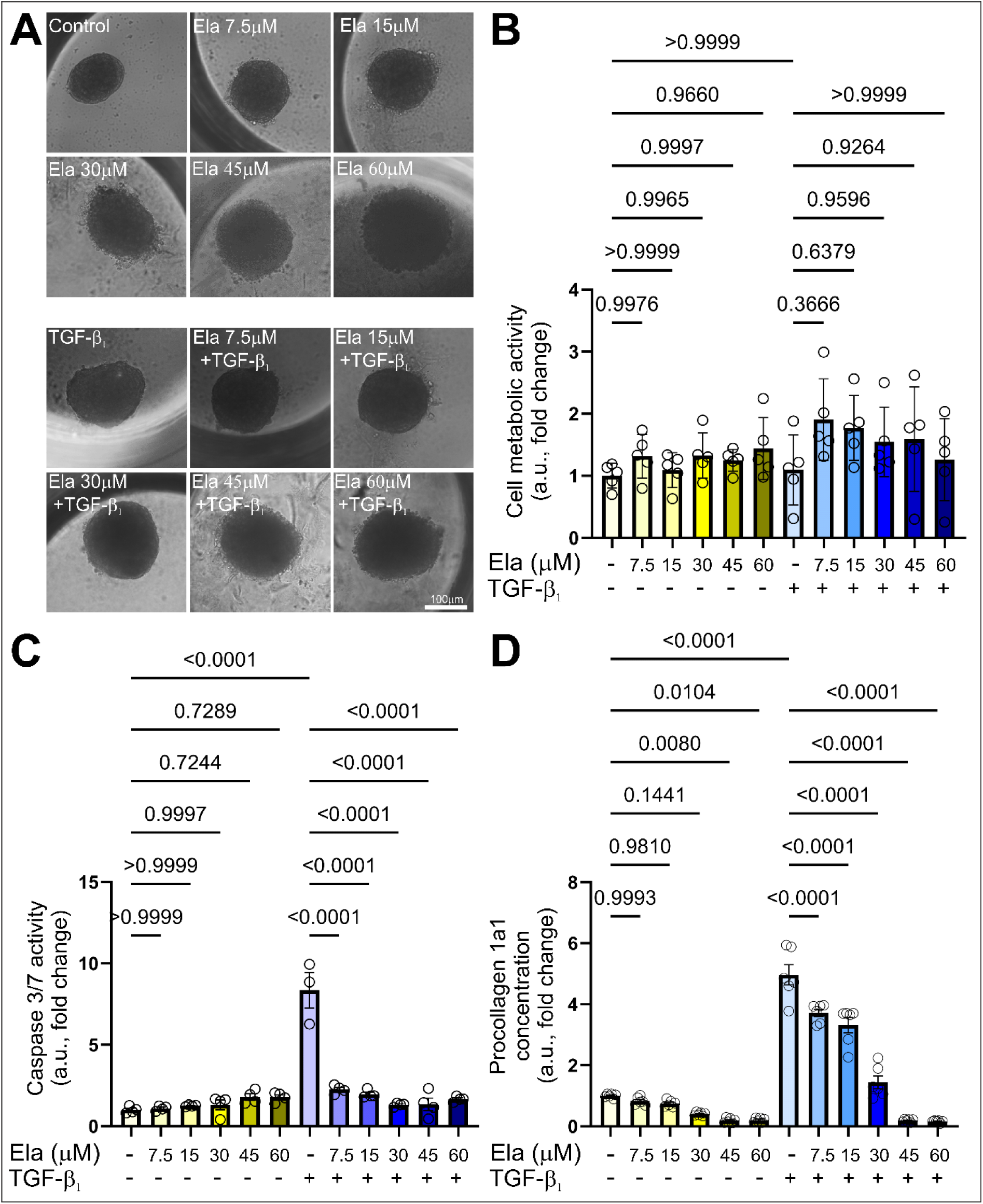
Elafibranor preserves viability and attenuates TGF-β₁-induced procollagen 1α1 secretion in 3D hCF spheroids. **(A)** Representative phase-contrast images of 3D spheroids treated with increasing concentrations of elafibranor (0-60 μM) in the absence or presence of TGF-β₁ (20 ng/ml) for 10 days. Scale bar = 100 μm. **(B)** Cell viability and **(C)** caspase-3/7 activity were measured in 3D hCF spheroids and expressed as arbitrary units (a.u.) in relation to control. **(D)** Procollagen 1α1 levels was determined by ELISA assay in collected supernatants after 3D hCF cultures. Data are presented as mean ± SD. Statistical significance was determined by one-way ANOVA followed by Tukey’s multiple comparisons test. p-values are indicated in each graph.

### 3.3. Elafibranor preserves viability and attenuates TGF-β_1_-induced caspase-3/7 activation in cardiac MTs

To investigate the effects of elafibranor in a physiologically relevant cardiac model, human cardiac MTs consisting of human induced pluripotent stem cell-derived cardiomyocytes (hiPSC-CMs) and human cardiac fibroblasts (hCFs) were employed as spontaneously contracting 3D tissues, enabling assessment of drug-induced changes in tissue viability and structural remodelling. Based on prior experiments, elafibranor was applied at 15 μM. Phase-contrast imaging showed that elafibranor did not alter MT morphology, irrespective of TGF-β₁ stimulation (Figure 4A). Consistently, resazurin-based assays indicated no reduction in cell viability following elafibranor treatment, either alone or in combination with TGF-β₁ (Figure 4B). TGF-β₁ significantly increased caspase-3/7 activity in cardiac MTs, whereas co-treatment with elafibranor reduced caspase-3/7 activity both under basal and TGF-β_1_-stimulated conditions (Figure 4C). Collectively, these findings indicate that elafibranor at 15 μM preserves MT morphology and viability while mitigating TGF-β₁-induced apoptotic signaling in human cardiac MTs.

**Figure 4.**
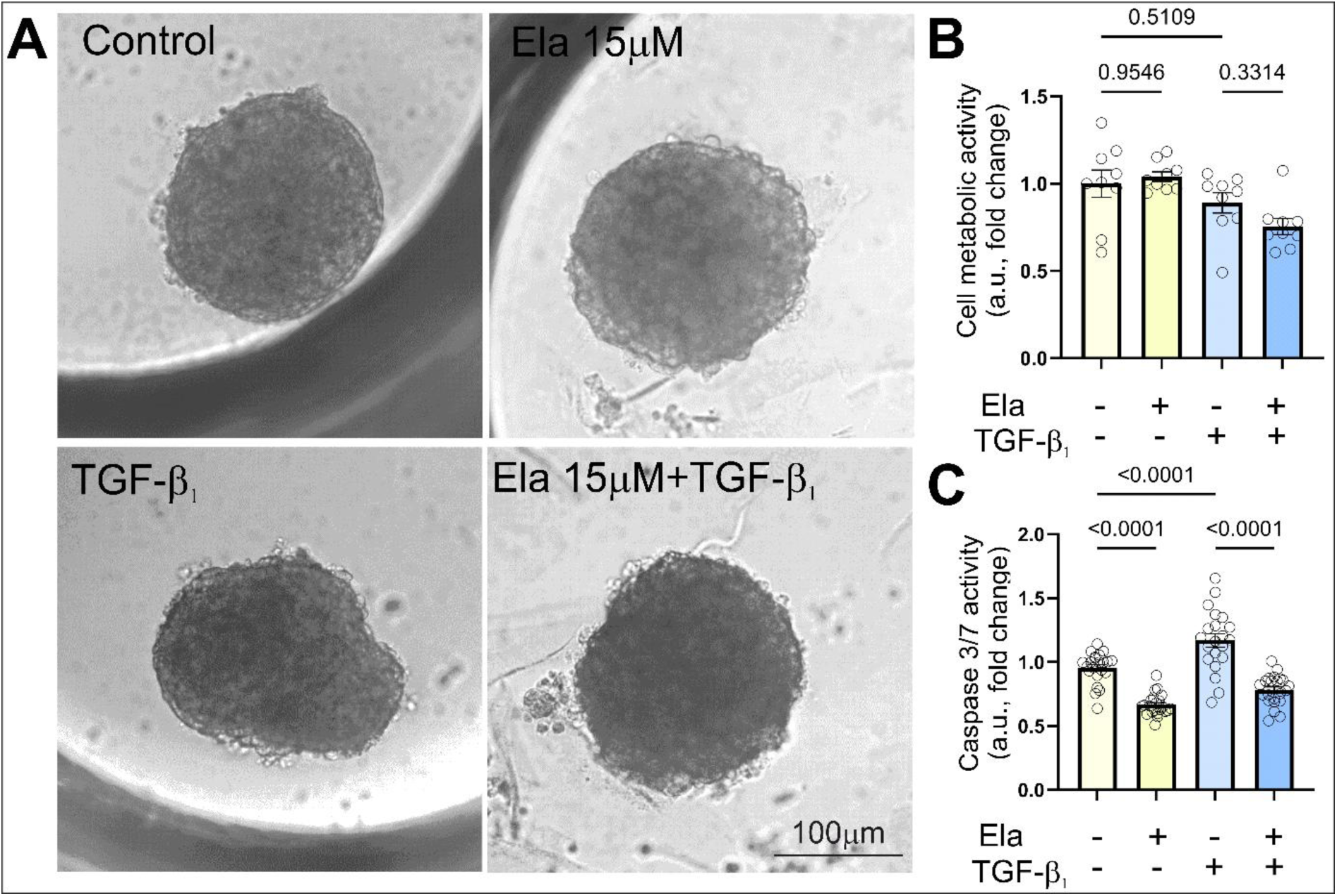
Elafibranor preserves morphology and viability while attenuating TGF-β_1_-induced caspase-3/7 activation in cardiac MTs. MTs was treated with elafibranor (15 μM) in the absence or presence of TGF-β_1_ (20 ng/ml) and cultured for 10 days. **(A)** Representative phase-contrast images of spherical MTs were presented. **(B)** Cell metabolic activity and **(C)** caspase 3/7 activity was measured in cultures of MTs at day 10. Data are presented as mean ± SD. Statistical significance was tested using the one-way ANOVA with Tukey’s post-hoc test. *p*-values are presented in each graph.

### 3.4. Elafibranor modulates the pro-fibrotic response of TGF-β₁-stimulated cardiac MTs

To investigate the molecular mechanisms underlying the effects of elafibranor and/or TGF-β_1_ on MT structure and function, global transcriptomic profiling was performed across experimental conditions using MTs treated with elafibranor at 15 μM in the absence or presence of TGF-β_1_. Differential expression analysis revealed condition-specific transcriptional signatures, as shown by Venn diagrams, hierarchical clustering of top DEGs and volcano plots (Figure 5A-C). In the elafibranor vs control comparison, upregulated genes included *TXNRD1, NQO1, UCP2, TXN,* and *LUCAT1*, indicating redox-regulatory, antioxidant, and mitochondrial/metabolic responses upon elafibranor treatment. The upregulation of canonical antioxidant/redox-associated genes such as *TXNRD1, NQO1,* and *TXN*, together with mitochondrial redox-related *UCP2*, is consistent with engagement of an NFE2L2/NRF2-associated cytoprotective program, although direct NRF2 activation was not assessed. In contrast, downregulated genes included *POSTN, ELN, MARCKS, CTGF*, and *MT1E*, consistent with reduced expression of genes associated with extracellular matrix remodeling, elastic fiber organization, profibrotic signaling, and stress-related programs. In the TGF-β₁ vs control comparison, upregulated genes included *PTK7, ENPP1, SDC1, ST6GAL2* and *HAPLN1*, whereas downregulated genes included *MYL3, HSPB8, MYL2, TNNC1* and *TNNI1*, supporting induction of cell-matrix interaction, proteoglycan/glycosaminoglycan biology, and extracellular matrix remodeling in response to TGF-β₁. In the TGF-β₁ + elafibranor vs TGF-β₁ comparison, upregulated genes included *AKR1B1, LAMA4, NMB, MYL3,* and *AKR1C2*, while downregulated genes included *ELN, MXRA5, HLA-B, MATN2,* and *ENPP1*. This profile suggests that elafibranor co-treatment shifts the TGF-β_1_-stimulated transcriptional response toward aldo-keto reductase/detoxification and stress-response programs, with additional basement membrane-associated, signaling-related, and partially contractile features, while reducing expression of selected TGF-β_1_-associated extracellular matrix/stromal genes. The ten top-ranked DEGs in each comparison are provided in Supplementary Tables S1-S3. Reactome GSEA performed on full ranked gene lists further demonstrated distinct pathway-level responses across treatment conditions (Figure 5D). Treatment of MTs by elafibranor alone was associated with enrichment of metabolic and mitochondrial pathways, including fatty acid metabolism and respiratory electron transport, whereas control MTs showed enrichment of extracellular matrix and collagen-related pathways. TGF-β₁ stimulation induced strong enrichment of cell-cycle-, DNA/RNA-processing-, protein synthesis-related pathways, and extracellular matrix organization in the full GSEA results. In the presence of TGF-β₁, elafibranor modulated TGF-β₁-associated transcriptional programs involving metabolic regulation, stress-response pathways, and extracellular matrix-associated processes.

**Figure 5.**
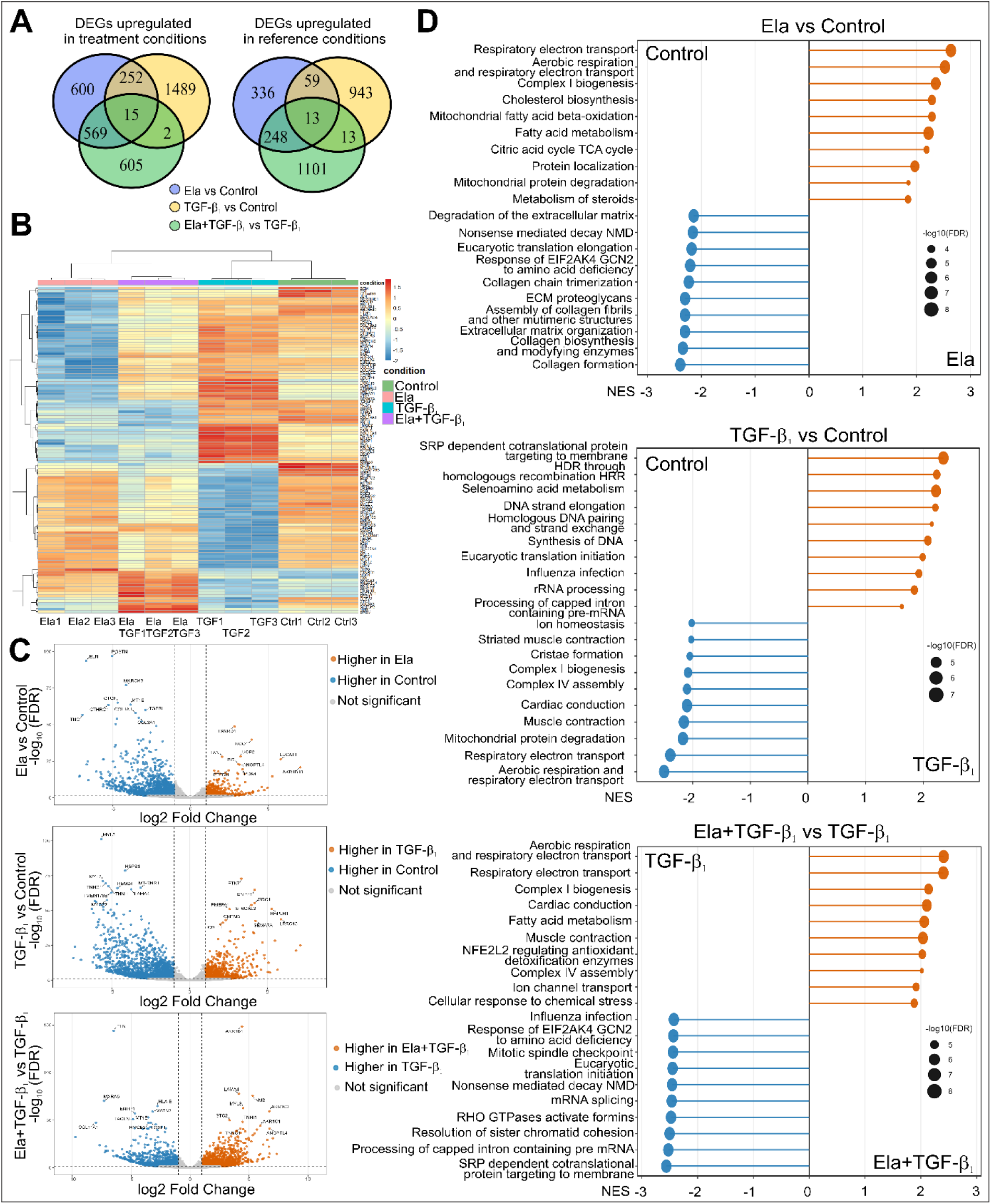
Elafibranor administered separately or together with TGF-β₁ modulates transcriptomic profiles of cardiac MTs. Transcriptional profiling and functional analysis of differential gene expression in MTs cultured in the absence or presence of TGF-β₁ (20 ng/ml) without or with elafibranor (15 μM) for 10 days. **(A)** Venn diagrams show overlaps of significant DEGs across the three oriented pairwise comparisons. Separate diagrams are shown for genes upregulated in the treatment condition and genes upregulated in the corresponding reference condition. **(B)** Heatmap of the union set of top DEGs, showing normalized expression across all samples with hierarchical clustering of genes and samples. **(C**) Volcano plots displaying DEGs identified using |log2 fold change| ≥ 1 and adjusted p-value < 0.05, with top genes labeled in each direction. **(D)** Reactome GSEA lollipop plots showing the top enriched pathways per comparison. NES indicates enrichment direction and strength, and dot size denotes −log10(FDR).

As a complementary DEG-based approach, Reactome ORA was performed separately for genes upregulated in each direction of the pairwise comparisons (Figure S1). ORA confirmed that elafibranor-upregulated DEGs were enriched in metabolic, mitochondrial, fatty acid metabolism, respiratory electron transport, and redox/antioxidant-related pathways, supporting a redox-adaptive transcriptional response consistent with NFE2L2/NRF2-associated signaling. In contrast, TGF-β₁-upregulated DEGs showed strong enrichment of extracellular matrix organization, collagen formation, elastic fibers formation, glycosaminoglycan-related pathways, and matrix remodeling, consistent with a profibrotic response. Combined TGF-β₁+ elafibranor treatment partially redirected this response toward redox-metabolic, detoxification/aldo-keto reductase-associated, mitochondrial/metabolic, and selected contractile or basement membrane-associated pathways, while several ECM-related pathways remained more prominent in TGF-β₁-treated MTs. Together, these results indicate that elafibranor remodels the transcriptional response of cardiac MTs by promoting redox-metabolic and cytoprotective programs, consistent with NFE2L2/NRF2-associated signaling, while reducing key extracellular matrix-related components of the TGF-β₁-driven profibrotic response.

To assess the relationship between transcriptomic changes and structural remodeling in MTs, immunocytochemical analyses were performed. Representative images of MTs cultured under control conditions or treated with elafibranor (15 μM), in the absence or presence of TGF-β₁, were stained for α-SMA, fibronectin, periostin, troponin T, and collagen (Masson’s trichrome) (Figure 6A). Control and elafibranor-treated cardiac MTs displayed low basal expression of fibrosis-associated markers and preserved sarcomeric organization, as indicated by troponin T staining. In contrast, TGF-β₁ stimulation markedly increased α-SMA, fibronectin, periostin, and collagen signals. Quantitative analyses demonstrated that elafibranor significantly attenuated TGF-β₁-induced upregulation of these fibrotic markers (Figure 6A-D, F). TGF-β₁ treatment also reduced troponin T expression, an effect that was partially reversed by elafibranor (Figure 6A, E). Consistently, TGF-β₁-induced secretion of procollagen 1α1 was significantly decreased upon elafibranor treatment (Figure 6G). Together, these findings indicate that elafibranor mitigates TGF-β₁-driven fibrotic remodeling of human cardiac MTs at the protein and tissue-structural level.

**Figure 6.**
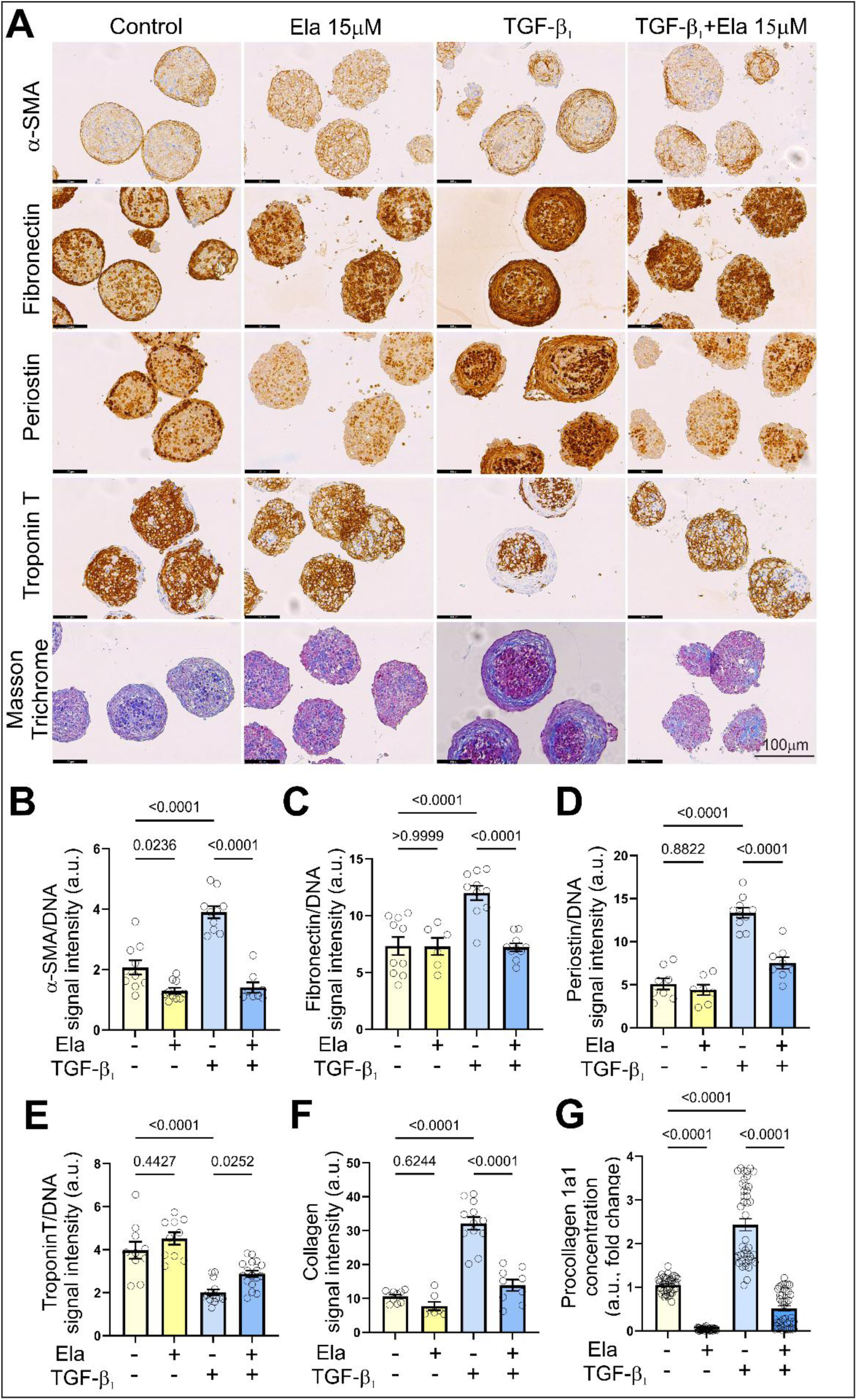
Elafibranor reduces TGF-β₁-induced fibrosis in human cardiac MTs. MTs treated with elafibranor (15 μM) in the absence or presence of TGF-β_1_ (20 ng/ml) cultured for 10 days were fixed, submerged in paraffin blocks, cut and stained. **(A)** Representative images of cardiac MTs stained for α-SMA, fibronectin, periostin, troponin T, and Masson’s trichrome. Scale bar = 100 μm. Quantification of **(B)** α-SMA, **(C)** fibronectin, **(D)** periostin and **(E)** Troponin T signal intensity normalized to DNA content. **(F)** Quantification of collagen signal from Masson’s trichrome staining determined in MTs. **(G)** Procollagen 1α1 concentration determined in supernatants from MTs cultures at day 10, presented in arbitrary units a.u.; *n* = 30. Data are presented as mean ± SD. Statistical significance was tested using the one-way ANOVA with Tukey’s post-hoc test. *p*-values are presented in each graph.

### 3.5. Elafibranor alters contractile properties and cellular bioenergetics in TGF-β₁-stimulated human cardiac MTs

To determine whether elafibranor-associated molecular and structural changes were accompanied by altered mechanical performance, cardiac MTs contractility was analyzed after 10 days of treatment using high-speed video recording and motion-tracking analysis [26,29,34]. TGF-β₁ stimulation changed the contractile profile of MTs by shortening contraction duration and relaxation time, reducing contraction amplitude, and increasing beating rate, without significantly affecting time-to-peak (Figure 7A-E). Elafibranor alone also affected contractile behavior, leading to shorter contraction duration, time-to-peak, and relaxation time, lower contraction amplitude, and a higher beating rate. When applied in the presence of TGF-β₁, elafibranor further accelerated contraction kinetics, as reflected by reduced contraction duration and time-to-peak, and significantly increased both contraction amplitude and beating rate, while relaxation time was not further changed compared with TGF-β₁ alone (Figure 7A-E). To evaluate whether these functional changes were associated with altered energetic status, adenine nucleotide ratios, total adenine nucleotide content, and NAD levels were quantified by an ultra-high performance liquid chromatography-diode array detection (UHPLC-DAD). ATP/AMP and ATP/ADP ratios remained statistically unchanged across the treatment groups, although both ratios tended to decrease in MTs exposed to combined TGF-β₁+ elafibranor treatment (Figure 7F-G). In contrast, and in line with the significant differences shown in panels H and I, elafibranor significantly reduced total adenine nucleotide (TAN) content and NAD levels in TGF-β₁-stimulated MTs compared with TGF-β₁ alone (Figure 7H-I). These findings indicate that elafibranor did not merely normalize TGF-β₁-induced dysfunction but instead reprogrammed MTs toward a distinct functional phenotype marked by accelerated beating, partial recovery of contractile strength, and decreased TAN and NAD pools under profibrotic conditions induced by TGF-β₁.

**Figure 7.**
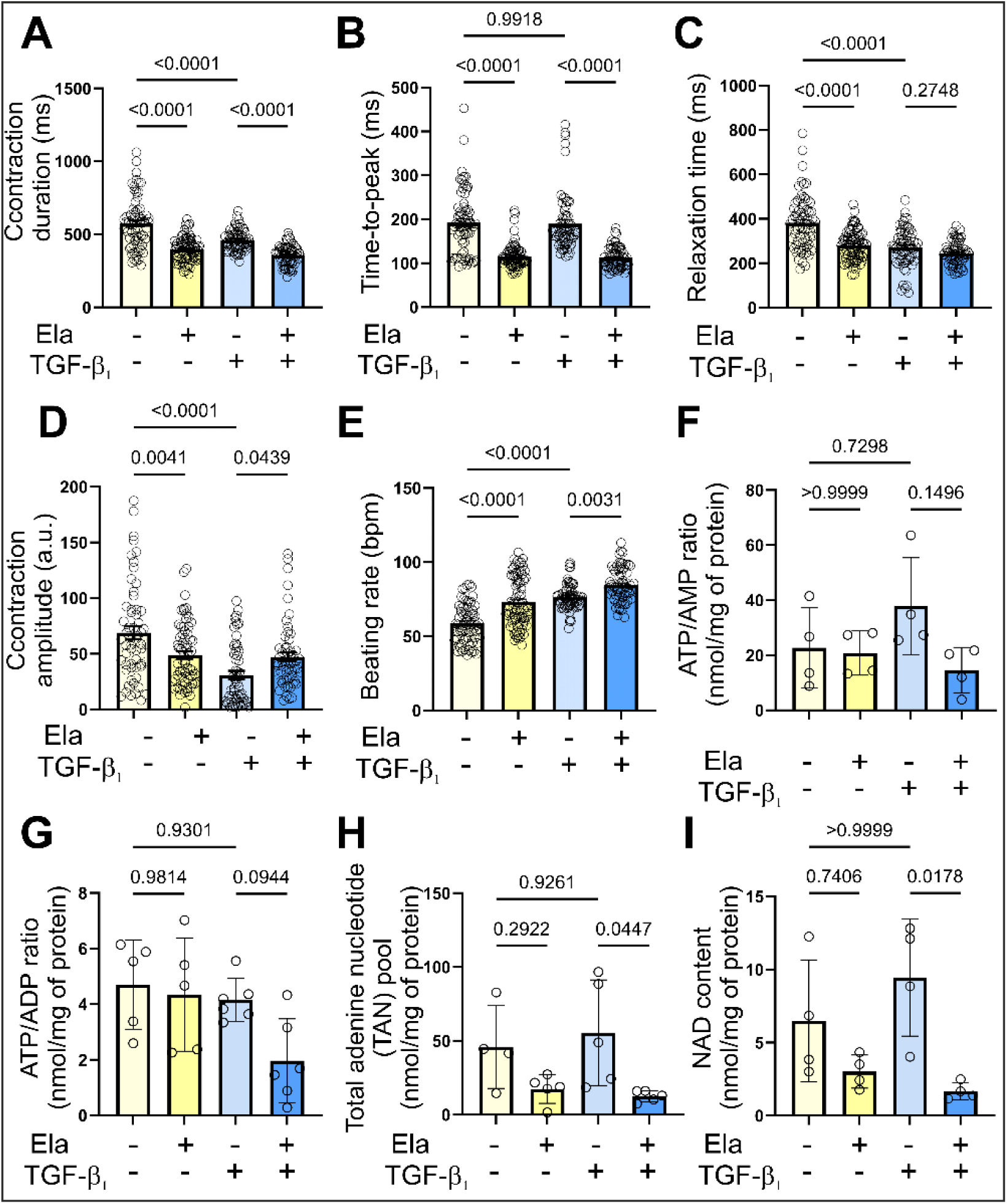
Elafibranor modifies contractility and nucleotide homeostasis in TGF-β_1_-stimulated cardiac MTs. Contraction of MTs were recorded for 15 s using an Axio Observer (Zeiss) microscope, and the MUSCLEMOTION macro together with Fiji ImageJ software was used to quantify contractile parameters: **(A)** contraction duration, **(B)** time-to-peak, **(C)** relaxation time, **(D)** contraction amplitude, and **(E)** beating rate. Each dot represents data from one MTs; at least 12 MTs were analyzed in each of three independent experiments. Nucleotide concentrations in MTs were determined using UHPLC-DAD method: **(F)** ATP/AMP ratio, **(G)** ATP/ADP ratio, **(H)** total adenine nucleotide (TAN) pool and **(I)** NAD content. Values were normalized to total protein content and are presented as mean ± SD (n ≥ 5). Statistical significance was assessed using one-way ANOVA followed by Tukey’s post hoc test; p values are indicated in the graphs.

### 3.6. Elafibranor modulates force generation, calcium handling, and mitochondrial respiration in TGF-β_1_-stimulated hiPSC-derived cardiomyocytes

Given the pronounced effects of elafibranor in human cardiac MT, consisting of hiPSC-CMs and hCFs mitigating TGF-β₁-induced fibrosis to assess the effects of hiPSC-CMs alone, we next investigated whether these functional changes were associated with alterations in force generation, calcium handling, and mitochondrial respiration in 2D cultures of hiPSC-CMs. Contraction force, frequency and calcium flux were analyzed using atomic force microscopy (AFM) combined with fluorescence imaging in untreated or TGF-β₁-treated cells in the absence or presence of elafibranor (15 μM). TGF-β₁ significantly increased contraction force and calcium flux while reducing beating rate compared with control cells. Elafibranor markedly attenuated the TGF-β₁-induced increases in force and calcium flux and partially restored beating rate (Figure 8A-D). Mitochondrial function was assessed by measuring oxygen consumption rates (OCR) using a Seahorse-based assay (Figure 8E). TGF-β₁ improved mitochondrial respiration, as evidenced by enhancement of basal respiration, maximal respiration, spare respiratory capacity, and ATP production. Elafibranor did not reduce the TGF-β₁-induced increase in basal or ATP-linked respiration; however, it significantly attenuated maximal respiration and spare respiratory capacity. Notably, elafibranor alone stimulated basal respiration and ATP production compared with control cells. Collectively, these results indicate that elafibranor modulates TGF-β₁-induced changes in contractile function, calcium handling, and mitochondrial bioenergetics in hiPSC-derived CMs.

**Figure 8.**
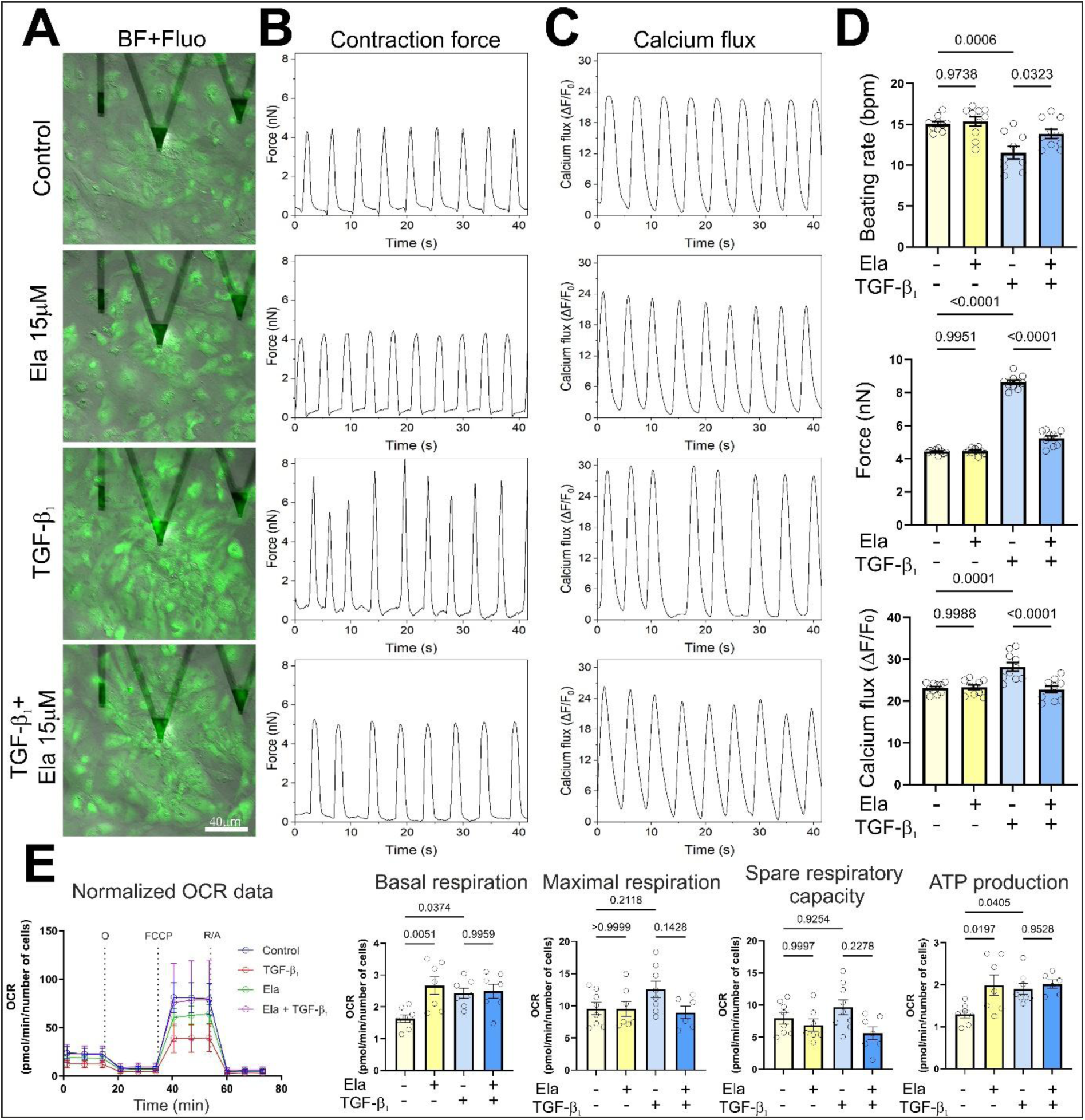
Elafibranor modulates contractile function, calcium handling, and mitochondrial bioenergetics in TGF-β₁-treated hiPSC-derived cardiomyocytes. Functional analysis of contractile properties and calcium flux in hiPSC-CMs treated with elafibranor (15 μM) in the absence or presence of TGF-β_1_ (10 ng/ml) using combined atomic force (AFM) and fluorescence microscopy. **(A)** Representative images (merged bright field (BF) and fluorescence (Fluo)) of hiPSC-CMs. Panels illustrate **(B)** cell contraction force determined by AFM and **(C)** calcium flux in hiPSC-CMs. **(D)** Quantitative analysis of the cell functional parameters: beating rate, force of contractions and calcium flux, n = 10. **(E)** Trajectories and OCR values were normalized to cell number for all tested conditions. Cellular bioenergetic changes was presented as basal respiration, maximal respiration, spare respiratory capacity and ATP-linked respiration. Data are presented as mean ± SD. Statistical significance was tested using the Kruskal-Wallis test with Dunn’s post-hoc test. p-values are presented in each graph.

## 4. Discussion

Elafibranor attenuates TGF-β_1_-induced profibrotic activation in human cardiac models, with effects linked to coordinated metabolic, redox, and functional remodeling. This is relevant because established ECM deposition and tissue stiffening remain difficult to reverse with standard cardiovascular therapies, whereas broad inhibition of central profibrotic pathways such as TGF-β_1_ may interfere with tissue repair and homeostatic signaling [4,35–39]. Elafibranor was used at 15 μM, consistent with previous non-cardiac studies in human hepatic NASH cultures, precision-cut liver slices treated with 10 μM elafibranor, and ALD-related hepatic or intestinal barrier assays using concentrations up to 30 μM [40–42]. At this concentration, elafibranor reduced α-SMA-positive myofibroblast formation, procollagen 1α1 secretion, and ECM-associated protein accumulation in 2D hCF cultures, 3D fibroblast spheroids, and cardiac MTs. These effects occurred without reduced viability and with lower TGF-β_1_-associated caspase-3/7 activity in 3D models, supporting an elafibranor-driven phenotypic effect rather than a response secondary to cytotoxicity. The distinct response profile observed in hiPSC-derived cardiomyocytes further suggests that elafibranor does not induce a uniform response across all cardiac cell types, but that its antifibrotic effect is primarily linked to modulation of fibroblast activation and ECM-associated changes.

The mechanistic interpretation is anchored in the pharmacology of elafibranor/GFT505 as a PPAR-α/δ agonist. In liver-centered models, elafibranor activates PPAR-linked fatty-acid metabolic programs and reduces inflammatory and ECM-related gene networks [23,42–44]. The relevance of PPAR signaling to fibroblast activation is supported by studies showing that PPAR-δ activation reduces proliferation, fibroblast-to-myofibroblast transition, and collagen synthesis in cardiac fibroblasts [45], whereas GW501516 attenuates TGF-β_1_-induced myofibroblast transition and ECM-marker expression in human bronchial fibroblasts through Smad2/p300/Sox9-related mechanisms [30]. PPAR-α may add a matrix-regulatory component, as elafibranor/GFT505 reduced Col1a1 and dermatopontin in liver fibrosis/NASH models, with dermatopontin repression linked to KLF6/TGF-β_1_ signaling [46]. Although receptor activation and these downstream mechanisms were not directly tested here, the convergence between the established metabolic pharmacology of elafibranor/GFT505 and the enrichment of lipid-metabolic, mitochondrial, and redox-related pathways in cardiac MTs supports a model in which PPAR-associated metabolic reprogramming may contribute to attenuation of the TGF-β_1_-induced profibrotic response.

Fibrotic activation is accompanied by metabolic reprogramming, and accumulating evidence indicates that these changes support myofibroblast differentiation, persistence, and ECM production [47–49]. This has been shown across fibrosis-relevant models: TGF-β_1_-induced fibroblast differentiation is associated with increased mitochondrial content and respiration; dermal fibroblast activation depends on glycolytic and glutamine reprogramming; and pulmonary fibrosis models implicate glycolysis, glutamine metabolism, and mTORC1/ATF4-dependent serine-glycine biosynthesis in myofibroblast differentiation and collagen production [50–55]. Combined activation of PPAR-β/δ and PPAR-γ with GW0742 and rosiglitazone reduces TGF-β_1_-induced collagen production while increasing peroxisomal biogenesis, lipid metabolism, and catalase-related redox control in primary human lung fibroblasts from control donors and patients with idiopathic pulmonary fibrosis [56]. Although elafibranor has been linked to metabolic and antifibrotic effects in hepatic and alcohol-related liver disease models, its impact on fibrosis-associated metabolism in human cardiac models has not been defined. We addressed this gap with complementary readouts. In hCFs, Seahorse analysis showed that elafibranor restored TGF-β_1_-impaired maximal respiration and spare respiratory capacity during fibroblast differentiation, in parallel with reduced α-SMA-positive myofibroblast formation and procollagen 1α1 secretion. In cardiac MTs, UHPLC-DAD quantified steady-state adenine nucleotide and NAD pools in whole multicellular tissues undergoing a TGF-β_1_-induced profibrotic response. Unlike Seahorse analysis, these measurements do not assess mitochondrial respiratory flux, but rather reflect the balance between synthesis, utilization, recycling, and consumption of nucleotides and redox cofactors. Therefore, the reduction in total adenine nucleotide and NAD pools should not be interpreted as impaired bioenergetics, particularly because ATP/AMP and ATP/ADP ratios were not significantly changed and adenylate energy charge was preserved in an auxiliary analysis (data not shown). Rather, these findings suggest altered nucleotide and redox-cofactor homeostasis in elafibranor-treated MTs, potentially involving changes in metabolite turnover or utilization. Because these measurements were performed in whole MTs, they cannot determine whether these changes originated primarily from cardiomyocytes, fibroblasts, or both cell populations. This interpretation was complemented by RNA-seq data pointing to broader metabolic reprogramming involving mitochondrial function, respiratory electron transport, lipid metabolism, and redox regulation. Enrichment of NFE2L2-regulated antioxidant programs, together with increased *TXNRD1, NQO1, TXN,* and *UCP2*, suggests a possible NRF2-related component. NRF2 activation has previously been associated with antifibrotic effects in systemic sclerosis/skin fibrosis models, TGF-β_1_-or IL-13-stimulated dermal fibroblasts, and TGF-β_1_-stimulated cardiac fibroblasts, where it reduced canonical TGF-β/Smad signaling, periostin expression, ECM production, or myofibroblast-associated contraction [57–59]. NRF2 activity was not directly measured, and bulk RNA-seq cannot assign this signal to a single cell population. Nevertheless, the respiratory, nucleotide/NAD, redox, and transcriptomic data connect elafibranor-induced metabolic reprogramming with attenuation of TGF-β_1_-induced profibrotic structural changes.

The contractile response should be interpreted through both MT structure and cardiomyocyte metabolism. In cardiac MTs, TGF-β_1_ increased ECM-associated protein accumulation, which can alter how cardiomyocytes generate and transmit force. By reducing ECM accumulation, elafibranor may have changed the mechanical environment of the tissue and contributed to partial recovery of contraction amplitude and changes in contraction kinetics. This structural effect is unlikely to fully explain the response, as elafibranor also altered adenine nucleotide and NAD pools at the whole-MT level and changed transcriptional programs related to mitochondrial function, ion handling, conduction, and muscle contraction. These changes are relevant because contraction depends on rapid energy transfer from mitochondrial ATP production to the contractile apparatus, supported by NAD-dependent redox reactions and the phosphocreatine system, rather than on total ATP levels alone [60–62]. The hiPSC-derived cardiomyocyte experiments were included to determine whether elafibranor also exerted cardiomyocyte-intrinsic effects beyond its fibroblast-associated antifibrotic activity. The distinct response observed in isolated hiPSC-derived cardiomyocytes indicates that elafibranor does not induce a uniform response across cardiac cell types: in hCFs, it restored TGF-β_1_-impaired respiratory reserve in parallel with reduced myofibroblast activation, whereas in cardiomyocytes it modulated a different TGF-β_1_-induced state characterized by altered mitochondrial respiration, calcium handling, and contraction. Thus, the tissue-level functional effect of elafibranor likely reflects both indirect effects related to reduced ECM accumulation and structural stress within MTs, and direct cardiomyocyte-level changes in mitochondrial respiration and calcium handling.

By combining hCFs, fibroblast spheroids, cardiac MTs, and hiPSC-derived cardiomyocytes, this study provides a multilayered human platform to examine how elafibranor affects profibrotic activation, ECM accumulation, metabolism, and contractile function. The same design also defines the scope of mechanistic interpretation. TGF-β_1_ stimulation captures an important profibrotic pathway but does not reproduce the inflammatory, vascular, mechanical, and metabolic complexity of cardiac fibrosis in vivo. Because elafibranor was co-administered with TGF-β_1_, our data address attenuation of a developing response rather than reversal of established fibrosis. We did not directly measure PPAR-α/δ activation, NRF2 activity, or TGF-β/Smad signaling; therefore, these pathways should be considered candidate mechanisms. Similarly, the respiratory, nucleotide/NAD, and transcriptomic data support elafibranor-associated metabolic remodeling, but do not resolve the relative contribution of glycolysis, fatty acid oxidation, metabolic fluxes, or cell-type-specific nucleotide/NAD changes. Future studies using receptor-selective or genetic approaches, cell-type-resolved omics, direct metabolic flux analyses, and models incorporating established fibrosis, mechanical load, and cardiometabolic stress should define how these pathways contribute to the cardiac effects of elafibranor.

## 5. Conclusion

Elafibranor attenuated the TGF-β_1_-induced profibrotic response in complementary human cardiac models. It reduced myofibroblast formation, procollagen 1α1 secretion, and ECM-associated protein accumulation without evidence of nonspecific cytotoxicity. These structural effects were accompanied by changes in mitochondrial respiration, nucleotide/NAD homeostasis, redox-associated transcriptional programs, cardiomyocyte calcium handling, and tissue-level contraction. To our knowledge, this is the first evidence connecting elafibranor with mitochondrial respiration, nucleotide/NAD homeostasis, and redox-associated transcriptional programs in human cardiac models of TGF-β_1_-induced profibrotic activation.

## List of abbreviations

α-SMA: α-smooth muscle actin
ADP: adenosine diphosphate
AFM: atomic force microscopy
AMP: adenosine monophosphate
ANOVA: analysis of variance
ATP: adenosine triphosphate
AU: arbitrary units
BF: bright field
BSA: bovine serum albumin
CVDs: cardiovascular diseases
DAB: 3,3′-diaminobenzidine
DEGs: differentially expressed genes
DMEM-HG: Dulbecco’s Modified Eagle Medium, high glucose
ECM: extracellular matrix
EDTA: ethylenediaminetetraacetic acid
ELA: elafibranor
ELISA: enzyme-linked immunosorbent assay
EtBr: ethidium bromide
FBS: fetal bovine serum
FCCP: carbonyl cyanide-p-trifluoromethoxyphenylhydrazone
FDA: fluorescein diacetate
FDR: false discovery rate
Fluo: fluorescence
GEO: Gene Expression Omnibus
GSEA: gene set enrichment analysis
hCF(s): human cardiac fibroblast(s)
HFpEF: heart failure with preserved ejection fraction
hiPSC-CM(s): human induced pluripotent stem cell-derived cardiomyocyte(s)
HRP: horseradish peroxidase
LDEV: lactate dehydrogenase-elevating virus
MM: maintenance medium
MT(s): microtissue(s)
MTT: 3-(4,5-dimethylthiazol-2-yl)-2,5-diphenyltetrazolium bromide
NAD: nicotinamide adenine dinucleotide
NASH: nonalcoholic steatohepatitis
NES: normalized enrichment score
NFE2L2/NRF2: nuclear factor erythroid 2-like 2 / nuclear factor erythroid 2-related factor 2
OCR: oxygen consumption rate
ORA: over-representation analysis
PBS: phosphate-buffered saline
PPAR(s): peroxisome proliferator-activated receptor(s)
RAAS: renin-angiotensin-aldosterone system
ROS: reactive oxygen species
SD: standard deviation
SEM: standard error of the mean
TAN: total adenine nucleotide pool
TGF-β / TGF-β_1_: transforming growth factor beta / beta 1
TMB: tetramethylbenzidine
UHPLC-DAD: ultra-high-performance liquid chromatography with diode array detection

## Declarations

### Ethics approval and consent to participate

Not applicable. All human cell types used in this study were obtained from commercial suppliers and no identifiable human participant data or newly collected human tissue were used.

### Consent for publication

Not applicable.

### Availability of data and materials

RNA sequencing data generated in this study will be deposited in a public gene expression repository (Gene Expression Omnibus, GEO) and will be made available upon publication. Accession numbers will be provided in the final version of the manuscript. All other data supporting the findings of this study will be available from the corresponding author upon reasonable request.

### Authors’ contributions

Conceptualization, M.P.;

Methodology, M.P., L.M., A.L., M.C., P.K., B.K.-Z., A.B., M.S., S.B.-W., D.W.;

Software, M.P., L.M.;

Validation, M.P.;

Formal Analysis, M.P.;

Investigation, M.P.;

Resources, M.P., P.B. and G.K.;

Data Curation, M.P.;

Writing-Original Draft Preparation, M.P.;

Writing-Review & Editing, L.M., A.L., M.C., B.K.-Z., A.B., M.S., S.B.-W., D.W., P.K., S.C., P.B., Z.M., O.D., J.C. G.K. and M.P.;

Visualization, M.P.;

Supervision, J.C. and G.K.;

Project Administration, M.P. and G.K.;

Funding Acquisition, M.P., P.B., Z.M., O.D. and G.K.

All authors have read and agreed to the published version of the manuscript.

## Acknowledgements

The authors thank the Center for Microscopy and Image Analysis of the University of Zurich for technical assistance in data acquisition. Graphical abstract was created in BioRender. Paw, M. (2026) https://BioRender.com/km01lrf.

## Declaration of generative AI and AI-assisted technologies in the manuscript preparation process

During the preparation of this work, the authors used ChatGPT by OpenAI to assist with language editing and text refinement. After using this tool, the authors reviewed and edited the content as needed and take full responsibility for the content of the published article. No generative AI or AI-assisted tools were used to create or alter figures, images, artwork, experimental data, or scientific results.

## Supplementary data

**Figure S1.**
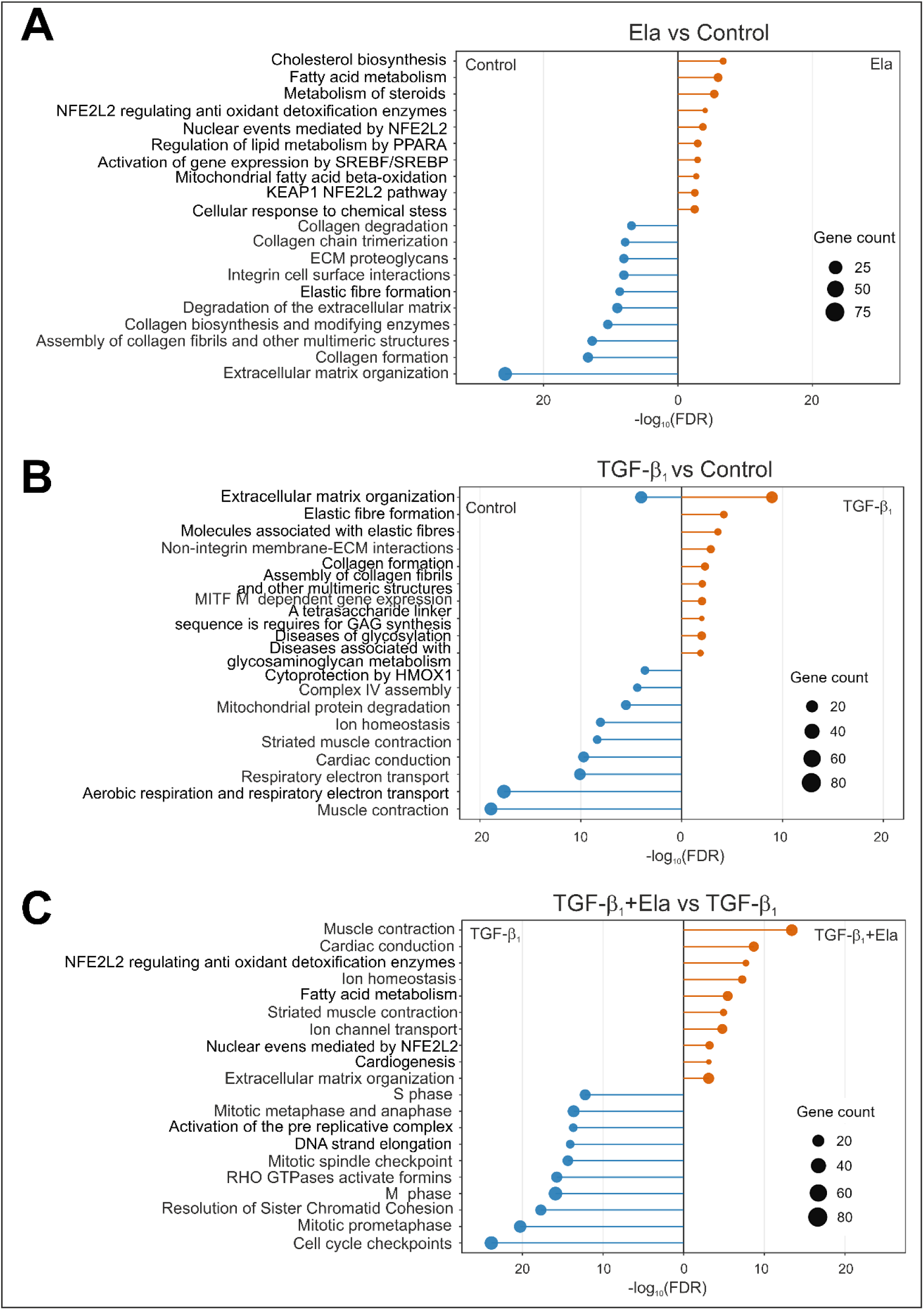
Reactome over-representation analysis of DEG sets across pairwise comparisons. Reactome ORA was performed separately for DEG sets upregulated in each direction of the indicated comparisons. DEGs were defined as |log₂FC| ≥ 1 and adjusted p-value < 0.05. Paired lollipop plots show significantly enriched Reactome pathways plotted toward the condition in which the corresponding DEG set was upregulated. The x-axis represents mirrored −log₁₀(FDR), and point size denotes the number of DEGs associated with each pathway. (A) In the elafibranor vs control comparison, elafibranor-upregulated DEGs were enriched in lipid, cholesterol, steroid, mitochondrial fatty acid β-oxidation, PPARα-related metabolism, and NFE2L2/NRF2-associated antioxidant detoxification pathways, whereas control-upregulated DEGs were enriched in extracellular matrix, collagen, elastic fibre, proteoglycan, and integrin-related pathways. (B) In the TGF-β₁ vs control comparison, TGF-β₁-upregulated DEGs showed enrichment of extracellular matrix and profibrotic pathways, including collagen formation, elastic fibre formation, non-integrin membrane-ECM interactions, and glycosaminoglycan/glycosylation-related processes, whereas control-upregulated DEGs were enriched in muscle contraction, cardiac conduction, respiratory electron transport, ion homeostasis, complex IV assembly and mitochondrial protein degradation pathways. (C) In the TGF-β₁+ elafibranor vs TGF-β₁ comparison, DEGs upregulated in the combined treatment were enriched in muscle contraction, cardiac conduction, ion homeostasis, fatty acid metabolism, NFE2L2/NRF2-associated antioxidant detoxification, cardiogenesis and extracellular matrix organization pathways. In contrast, DEGs upregulated in TGF-β₁-treated MTs were enriched in cell-cycle, S phase, mitotic checkpoint, DNA replication, sister chromatid cohesion and RHO GTPase/formin-associated pathways.

**Table 1.**
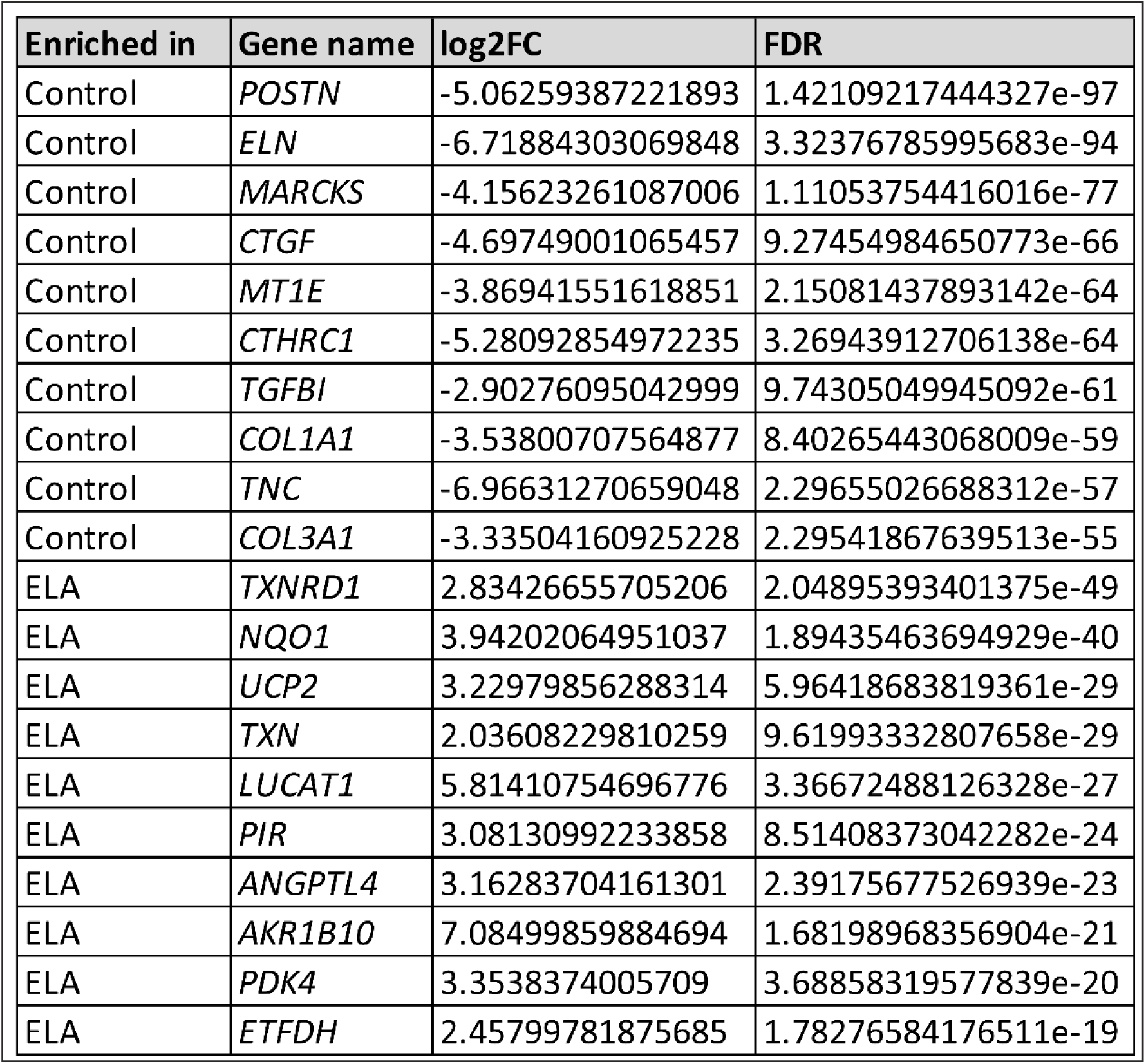
List of top differentially expressed genes in the elafibranor (ELA) vs control comparison. Differentially expressed genes (DEGs) were defined as genes with FDR < 0.05 and |log₂FC| ≥ 1. The table shows the top 10 genes upregulated in control and the top 10 genes upregulated in elafibranor-treated MTs, selected from the DEG list using a predefined ranking procedure. Positive log₂FC values indicate higher expression in elafibranor-treated MTs, whereas negative log₂FC values indicate higher expression in control.

| Enriched in | Gene name | log2FC | FDR |
| --- | --- | --- | --- |
| Control | *POSTN* | -5.06259387221893 | 1.42109217444327e-97 |
| Control | *ELN* | -6.71884303069848 | 3.32376785995683e-94 |
| Control | *MARCKS* | -4.15623261087006 | 1.11053754416016e-77 |
| Control | *CTGF* | -4.69749001065457 | 9.27454984650773e-66 |
| Control | *MT1E* | -3.86941551618851 | 2.15081437893142e-64 |
| Control | *CTHRC1* | -5.28092854972235 | 3.26943912706138e-64 |
| Control | *TGFBI* | -2.90276095042999 | 9.74305049945092e-61 |
| Control | *COL1A1* | -3.53800707564877 | 8.40265443068009e-59 |
| Control | *TNC* | -6.96631270659048 | 2.29655026688312e-57 |
| Control | *COL3A1* | -3.33504160925228 | 2.29541867639513e-55 |
| ELA | *TXNRD1* | 2.83426655705206 | 2.04895393401375e-49 |
| ELA | *NQO1* | 3.94202064951037 | 1.89435463694929e-40 |
| ELA | *UCP2* | 3.22979856288314 | 5.96418683819361e-29 |
| ELA | *TXN* | 2.03608229810259 | 9.61993332807658e-29 |
| ELA | *LUCAT1* | 5.81410754696776 | 3.36672488126328e-27 |
| ELA | *PIR* | 3.08130992233858 | 8.51408373042282e-24 |
| ELA | *ANGPTL4* | 3.16283704161301 | 2.39175677526939e-23 |
| ELA | *AKR1B10* | 7.08499859884694 | 1.68198968356904e-21 |
| ELA | *PDK4* | 3.3538374005709 | 3.68858319577839e-20 |
| ELA | *ETFDH* | 2.45799781875685 | 1.78276584176511e-19 |

**Table 2.** List of top differentially expressed genes in the TGF-β₁ vs control comparison. Differentially expressed genes (DEGs) were defined as genes with FDR < 0.05 and |log₂FC| ≥ 1. The table shows the top 10 genes upregulated in control and the top 10 genes upregulated in TGF-β₁-treated MTs, selected from the DEG list using a predefined ranking procedure. Positive log₂FC values indicate higher expression in TGF-β₁-treated MTs, whereas negative log₂FC values indicate higher expression in control.

| Enriched in | Gene name | log2FC | FDR |
| --- | --- | --- | --- |
| Control | *MYL3* | -5.6742321511674 | 6.76136986104236e-102 |
| Control | *HSPB8* | -4.14396404607516 | 2.91922877698559e-79 |
| Control | *MYL2* | -5.57355821074484 | 7.59165876581933e-72 |
| Control | *TNNC1* | -5.39825013409831 | 3.9267435607697e-70 |
| Control | *TNNI1* | -5.21477372054359 | 2.81234689743524e-68 |
| Control | *MT-RNR1* | -3.15075165758278 | 3.11529886956055e-67 |
| Control | *RBM24* | -4.64045887611901 | 6.55999039182358e-67 |
| Control | *LAMA4* | -3.75655586625305 | 6.29457508943034e-66 |
| Control | *TMEM176B* | -5.02162066071816 | 1.65018341873918e-63 |
| Control | *MYOZ2* | -5.30921912091656 | 2.7273897355702e-58 |
| TGF-$\beta_1$ | *PTK7* | 3.32803584385169 | 1.47798386043166e-73 |
| TGF-$\beta_1$ | *ENPP1* | 4.17769030808612 | 1.14091390027126e-65 |
| TGF-$\beta_1$ | *SDC1* | 4.17560677747325 | 4.24223206632889e-56 |
| TGF-$\beta_1$ | *ST6GAL2* | 3.97050475662376 | 1.0712951099381e-54 |
| TGF-$\beta_1$ | *HAPLN1* | 5.28281627647706 | 7.58202056228287e-52 |
| TGF-$\beta_1$ | *PMEPA1* | 2.56508292830819 | 9.78395192580278e-52 |
| TGF-$\beta_1$ | *LRRC15* | 5.87391624578281 | 1.78286643086965e-44 |
| TGF-$\beta_1$ | *CMTM3* | 2.3231035093485 | 5.80002878896648e-44 |
| TGF-$\beta_1$ | *SEMA7A* | 4.2372976336584 | 2.18777552214423e-43 |
| TGF-$\beta_1$ | *CDH11* | 2.13398569781797 | 5.40253260511048e-42 |

**Table 3.** List of top differentially expressed genes in the elafibranor (ELA) + TGF-β₁ vs TGF-β₁ comparison. Differentially expressed genes (DEGs) were defined as genes with FDR < 0.05 and |log₂FC| ≥ 1. The table shows the top 10 genes upregulated in TGF-β₁-treated MTs and the top 10 genes upregulated in MTs treated with ELA + TGF-β₁, selected from the DEG list using a predefined ranking procedure. Positive log₂FC values indicate higher expression in MTs treated with Ela + TGF-β₁, whereas negative log₂FC values indicate higher expression in TGF-β₁-treated MTs.

| Enriched in | Gene name | log2FC | FDR |
| --- | --- | --- | --- |
| TGF-$\beta_1$ | *ELN* | -6.48688060271927 | 1.16915210577081e-144 |
| TGF-$\beta_1$ | *MXRA5* | -7.30916524791498 | 7.08981950247325e-71 |
| TGF-$\beta_1$ | *HLA-B* | -2.74971194110169 | 1.26749993345745e-65 |
| TGF-$\beta_1$ | *MATN2* | -3.19246992388374 | 1.00609871822661e-59 |
| TGF-$\beta_1$ | *ENPP1* | -4.67612411939045 | 5.82221296449069e-58 |
| TGF-$\beta_1$ | *MT1E* | -3.54887069248333 | 1.40516658377722e-53 |
| TGF-$\beta_1$ | *TAGLN* | -4.8423488317734 | 1.53980122304982e-51 |
| TGF-$\beta_1$ | *COL11A1* | -7.98647157546631 | 3.95707922548347e-48 |
| TGF-$\beta_1$ | *HMGB2* | -3.54558440668677 | 4.61838114171094e-48 |
| TGF-$\beta_1$ | *LOXL1* | -3.33194321611303 | 8.71399508937439e-48 |
| ELA+ TGF-$\beta_1$ | *AKR1B1* | 4.40155546799057 | 4.06592081076487e-149 |
| ELA+ TGF-$\beta_1$ | *LAMA4* | 4.15325994443132 | 2.18991913838537e-78 |
| ELA+ TGF-$\beta_1$ | *NMB* | 5.30465458423155 | 1.28157848339648e-76 |
| ELA+ TGF-$\beta_1$ | *MYL3* | 4.51024634628406 | 2.12886835170813e-63 |
| ELA+ TGF-$\beta_1$ | *AKR1C2* | 6.71673932026951 | 3.96504986537609e-60 |
| ELA+ TGF-$\beta_1$ | *STC2* | 3.33325496487584 | 9.51109060730103e-51 |
| ELA+ TGF-$\beta_1$ | *TNNI1* | 4.45178126429389 | 5.24210085621984e-49 |
| ELA+ TGF-$\beta_1$ | *AKR1C1* | 5.95715540799631 | 8.61189721465161e-45 |
| ELA+ TGF-$\beta_1$ | *ANGPTL4* | 6.72908473281287 | 3.92669891284361e-42 |
| ELA+ TGF-$\beta_1$ | *TNNC1* | 4.26836053451261 | 4.60617646629787e-42 |

